# Archaeal signalling networks - new insights into the structure and function of histidine kinases and response regulators of the methanogenic archaeon *Methanosarcina acetivorans*

**DOI:** 10.1101/2024.10.04.616672

**Authors:** Nora FK Georgiev, Anne L Andersson, Zoe Ruppe, Loriana Kattwinkel, Nicole Frankenberg-Dinkel

## Abstract

The methanogenic archaeon Methanosarcina *acetivorans* has one of the largest known archaeal genomes. With 53 histidine kinases (HK), it also has the largest set of signal transduction systems. To gain insight into the hitherto not very well understood signal transduction in archaea and *M. acetivorans* in particular, we have categorized the predicted HK into four types based on their H-box using an *in silico* analysis. Representatives of three types were recombinantly produced in *Escherichia coli* and purified by affinity chromatography. All investigated kinases showed ATP binding and hydrolysis. The MA_type 2 kinase, which lacks the classical H-box, showed no autokinase activity. Furthermore, we could show that *M. acetivorans* possesses an above-average number of response regulators (RR), many of them consisting of only a REC domain (REC-only). Using the hybrid kinase MA4377 as an example we show that both intra-and intermolecular transphosphorylation to REC domains occur. These experiments are furthermore indicative of complex phosphorelay systems in *M. acetivorans* and suggest that REC-only proteins act as a central hub in signal transduction in *M. acetivorans*.

## Introduction

Sensing and responding to environmental stimuli are key abilities of microorganisms to adapt to their surroundings. Organisms detect stimuli through receptors, and in the process of signal transduction, they are transmitted through the cell in a series of events that ultimately lead to a cellular response (Stock *et al*., 2000). Signal transduction systems are known in all three domains of life and at different levels of complexity. In prokaryotes, it comprises one-component systems (OCS), two-component systems (TCS), and multistep phosphorelay systems. The less complex OCSs are believed to be evolutionary ancestors of TCSs. They are more widely distributed in prokaryotes, and their structure is based on fused input-output components (Ulrich *et al*., 2005). The classical TCS consists of a histidine sensor kinase (HK) and a response regulator (RR). The HK is usually membrane bound and harbours two main components, a sensor input domain and a transmitter domain, while the RR is usually a cytoplasmic protein with receiver domain (REC) and an output domain (Wolanin *et al*., 2002). The catalytic core of the HK consists of a dimerisation and histidine-phosphotransfer (DHp) domain and the catalytic and ATP-binding (CA) domain (Stock *et al*., 2000; Zschiedrich *et al*., 2016). Through signal input, a histidine residue within the so-called H-box of the DHp domain is phosphorylated by the γ-phosphoryl group of ATP. In the next step, this phosphoryl group is transferred to the cognate RR. To transfer phosphate from the HK to the RR, the RR docks onto the DHp domain and causes the phosphorylation of an aspartate residue, using the phosphohistidine of the HK as a substrate (Zschiedrich *et al*., 2016). Phosphorylation of the RR leads to the activation of its output domain, which triggers a cellular response. While a TCS consists of a sensor HK and a RR, there is also a more complex variant, the phosphorelay system. Here, a hybrid HK with a bound receiver domain transfers the phosphate to a histidine phosphotransfer (HPt) domain and then to a RR with output domain. The phosphorylation cascade of such systems proceeds through a four-step phosphorelay (His1-Asp1-His2-Asp2). Phosphorelay systems are found in prokaryotes and eukaryotes. While it was originally assumed that they occur most frequently in eukaryotes, it is now known that they are also very common in Bacteria and Archaea (Zhang & Shi, 2005). Although we know a lot about the signal transduction in Bacteria and Eukarya, the knowledge about archaeal signal transduction is still limited. It is thought that signal transduction systems evolved in bacteria after their separation from the last common ancestor, and Archaea and Eukarya received them through multiple horizontal gene transfer events (Brown et al. 2000). Thus far, only a few signal transduction systems of Archaea have been experimentally studied in detail.

To date, experimental data of three archaeal signal transduction are available. The chemotaxis protein CheA of *Halobacterium salinarum* was among the first HK identified in Archaea and has a function in taxis control as its bacterial counterpart (Rudolph & Oesterhelt, 1995). In addition, two TCS from methanogenic Archaea have been studied which are involved in the regulation of methanogenesis (FilI/FilR TCS from *Methanosaeta harundinacea*) and temperature sensing (LtrK/LtrR from *Methanococcoides burtonii*), respectively. All three systems work like typical bacterial TCS, wherein a HK is autophosphorylated, subsequently transferring the phosphoryl group to the REC domain of a cognate RR. The FilI/FilR systems involves one RR with putative DNA binding domain and a second REC-only RR with unknown function. While the LtrK/LtrR system uses a typical HTH-domain fused to the REC domain (Li *et al*., 2014; Najnin *et al*., 2016). In general, it is not yet fully understood how these rather bacterial-like components interact with the archaeal transcription machinery.

As these studies might not be representative of the whole domain of Archaea and unique archaeal systems might exist, we designed a study to investigate signal transduction in the methanogenic archaeon *M. acetivorans*. With 53 sensor kinases and 15 RRs, this archaeon possesses one of the largest signal transduction networks within the Archaea (Galperin *et al*., 2018). Using bioinformatics, we grouped the HK into four types based on the HK domain and investigated representative members of each type. Except one type, all investigated HK are true autokinases. Finally, we provide biochemical evidence for the existence of a (multiple) phosphorelay systems in *M. acetivorans*.

## Results

### Distribution of histidine kinases and response regulators in the genome of *M. acetivorans*

To gain a general overview of kinases and RRs in *M. acetivorans*, data of putative signal transduction systems from databases (MIST 4.0, P2CS) and literature were analysed (Galperin *et al*., 2018; Gumerov *et al*., 2023; Ortet *et al*., 2015). All examined sources showed that the genome of *M. acetivorans* encodes 53 putative HKs with characteristic cytoplasmic DHp (**D**imerisation and **H**istidine **p**hosphotransfer) and CA domains (**C**atalytic **A**TP-binding) and 15 putative RRs with REC domain (**REC**eiver). The HK include different sensing domains, mainly PAS (**P**er-**A**rnt-**S**IM), PAC (**P**AS-**A**ssociated **C**-terminal motif) and GAF domains (c**G**MP-specific phosphodiesterases, **a**denylyl cyclases and **F**hlA), but also CHASE4 (**C**yclases/**H**istidine kinases **A**ssociated **S**ensory **E**xtracellular), novel MEDS (**ME**thanogen/methylotroph **D**cmR **S**ensory) and PocR (sensory domain found in PocR, with novel variant of the PAS-like fold) domains. Among the 53 HK, eight are membrane bound while all other HK are cytosolic proteins. Furthermore, two proteins have an unusual domain organisation with a N-terminal REC domain. Three HKs are predicted to be hybrid histidine kinases (HHKs) with bound REC domains at their C-terminus. The ratio of HK to RR is around 3:1, which is rather unusual if compared to bacteria wherein the ratio is 1:1 (Galperin *et al*., 2018; Krell, 2018). Looking into their genome localisation, it can be distinguished between orphan kinases and kinases with genomic proximity to a RR (Figure 1A). The genes of seven putative HK are encoded in operons with one or more RR, while most of the HK are orphan.

**Figure 1:**
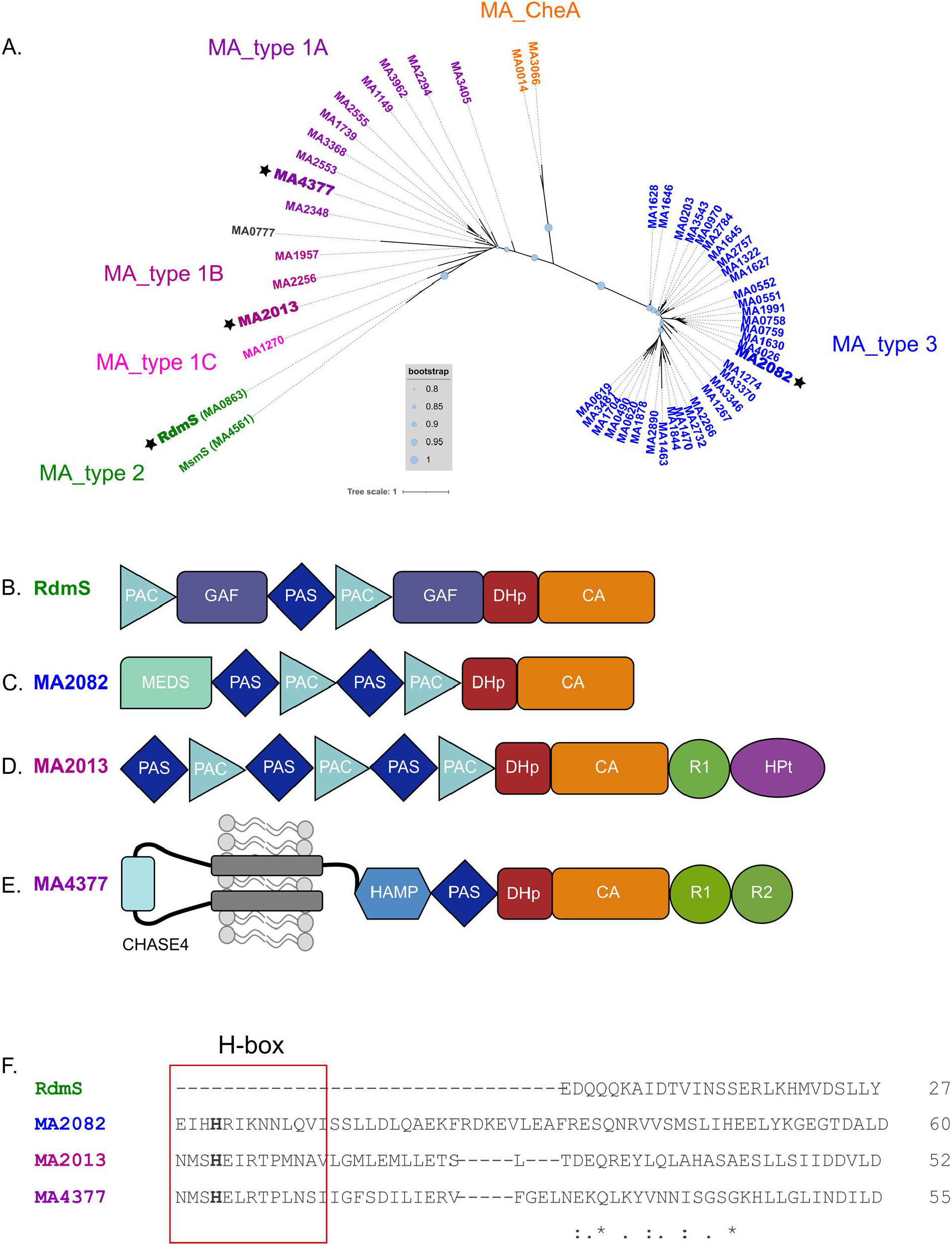
Unrooted phylogenetic tree depicting the HK domain of 53 putative HKs from *M. acetivorans* and schematic representation of the domain structure of four selected proteins. (**A**.) Phylogenetic analysis of the HK domain from *M. acetivorans*. The maximum-likelihood phylogenetic tree was constructed using PhyML 3.0 (Guindon *et al*., 2010), bootstrap values were calculated using the online tool BOOSTER (Lemoine *et al*., 2018). Bootstrap values are displayed as light blue circles in different sizes. To generate the tree, only the HK domain of the HK were considered. Branches are coloured according to which HK type is present. Accession numbers of the used sequences are found in Supplement Table S4. (**B**.) Domain structure of the protein RdmS, a cytosolic HK lacking the H-box. (**C**.) Domain structure of the protein MA2082, a cytosolic histidine kinase. (**D**.) Domain structure of the protein MA2013, a cytosolic hybrid histidine kinase with C-terminal REC and Hpt domain. (**E**.) Domain structure of the protein MA4377, a membrane bound hybrid histidine kinase with two bound REC domains. The domain structures of the four proteins were analysed using the programs SMART (Letunic *et al*., 2020) and InterPro (Paysan-Lafosse *et al*., 2023). PAS domain (**P**er-**A**rnt-**S**IM), PAC domain (**P**AS-**A**ssociated **C**-terminal motif), GAF domain (c**G**MP-specific phosphodiesterases, **a**denylyl cyclases and **F**hlA), CHASE4 domain (**C**yclases/**H**istidine kinases **A**ssociated **S**ensory **E**xtracellular), HAMP domain (**H**istidine kinases, **A**denylate cyclases, **M**ethyl accepting proteins and **P**hosphatases), MEDS domain (**ME**thanogen/methylotroph **D**cmR **S**ensory), DHp domain (**D**imerisation and **H**istidine **p**hosphotransfer), CA domain (**C**atalytic **A**TP-binding), R (**R**eceiver domain), HPt domain (**H**istidine **P**hospho**t**ransfer). (**F**.) Alignment of HK domain of all four putative HK, H-box region is highlighted with a red box.

Bacterial HKs can be categorised into five different groups (Type I, Type II, Type III, Type IV, CheA) according to the structure of their phosphorylation site (H-box) (Kim & Forst, 2001). An amino acid sequence alignment and a phylogenetic tree of the HK domain of all 53 kinases of *M. acetivorans* showed that they group into four distinct groups depending on the amino acid sequence of the H-box region (Figure 1A, Supplement Figure S2). As only one of these groups belongs to the known bacterial kinase types, new categories were designated for *M. acetivorans* (Figure 1A). They were named MA_type 1A/B/C, MA_type 2, MA_CheA and MA_type3. MA_type 1 is similar to the Type I HK of bacteria (Kim & Forst, 2001). As the amino acid arrangement differs within the group, the subgroups A, B and C were created. MA_type 2 kinases are characterised by the absence of an H-box and MA_type 3 kinases cannot be grouped with a known type of bacterial kinases. MA_CheA are highly similar to bacterial CheA proteins.

The analyses of the putative RRs revealed that eleven of them are so-called REC-only proteins and only contain a REC domain, which is in stark contrast to bacterial systems, wherein REC domains are most often connected to output domains. Four of the RRs show the characteristic bacterial domain organisation with a REC domain and a fused output domain. Two of them have a fused CheB domain, one an HTH domain and one a MetOD1 domain, which is a specific domain for methanogens with yet unknown function (Galperin *et al*., 2018). An amino acid sequence alignment of the REC domain of all RRs showed an overall structural similarity (Supplement Figure S3).

### Biochemical analysis of four putative histidine kinases of *M. acetivorans*

In order to investigate the biochemical commonalities or differences and to get more functional insight into archaeal signal transduction, one member of each newly defined type was chosen for further investigation. In addition, for MA_ type 1, we picked a membrane bound (MA4377, MA_type 1A) and a cytoplasmic representative (MA2013, MA_type 1B) (Figure 1B-E). Both kinases are encoded next to a REC-only RR in the genome. RdmS (MA0863) is an orphan cytoplasmic kinase lacking the H-box (Fiege & Frankenberg-Dinkel, 2019), and a representative of MA_type 2. MA2082 is an orphan, cytoplasmic histidine kinase possessing a methanogen-specific MEDS domain and is grouped as a MA_type 3 kinase.

### All examined kinases bind and hydrolyse ATP

The function of HKs relies on the binding of ATP to the ATP binding site in the CA domain and its hydrolysis in the DHp domain, which results in the transfer of the γ-phosphate to the conserved His residue situated in the H-box. In order to test ATP binding and hydrolysis, all four HK were recombinantly produced in *E. coli* and purified using affinity chromatography.

ATP binding was tested employing the fluorescent ATP-derivative 2’-(3’)-O-(2,4,6-trinitrophenyl)-adenosin 5’-triphosphat (TNP-ATP). For all tested kinases, the fluorescence signal clearly increased upon incubation with TNP-ATP. This was also true for a MA4377 variant lacking the membrane domain (PKR1R2). All kinases displayed affinity towards ATP with K_d_ values in the micromolar range with MA4377 and its variants having the highest affinity (K_d_ ∼ 11 µM) (Figure 2A, Supplement Figure S4). We next asked, whether the kinases are also able to hydrolyse the bound ATP. Using the commercial ADP-Glo^™^ Kinase Assay (Promega), all four kinases were not only shown to bind but also to hydrolyse ATP (Figure 2B). Therefore, the next step was to test their autokinase activity via autophosphorylation assays.

**Figure 2:**
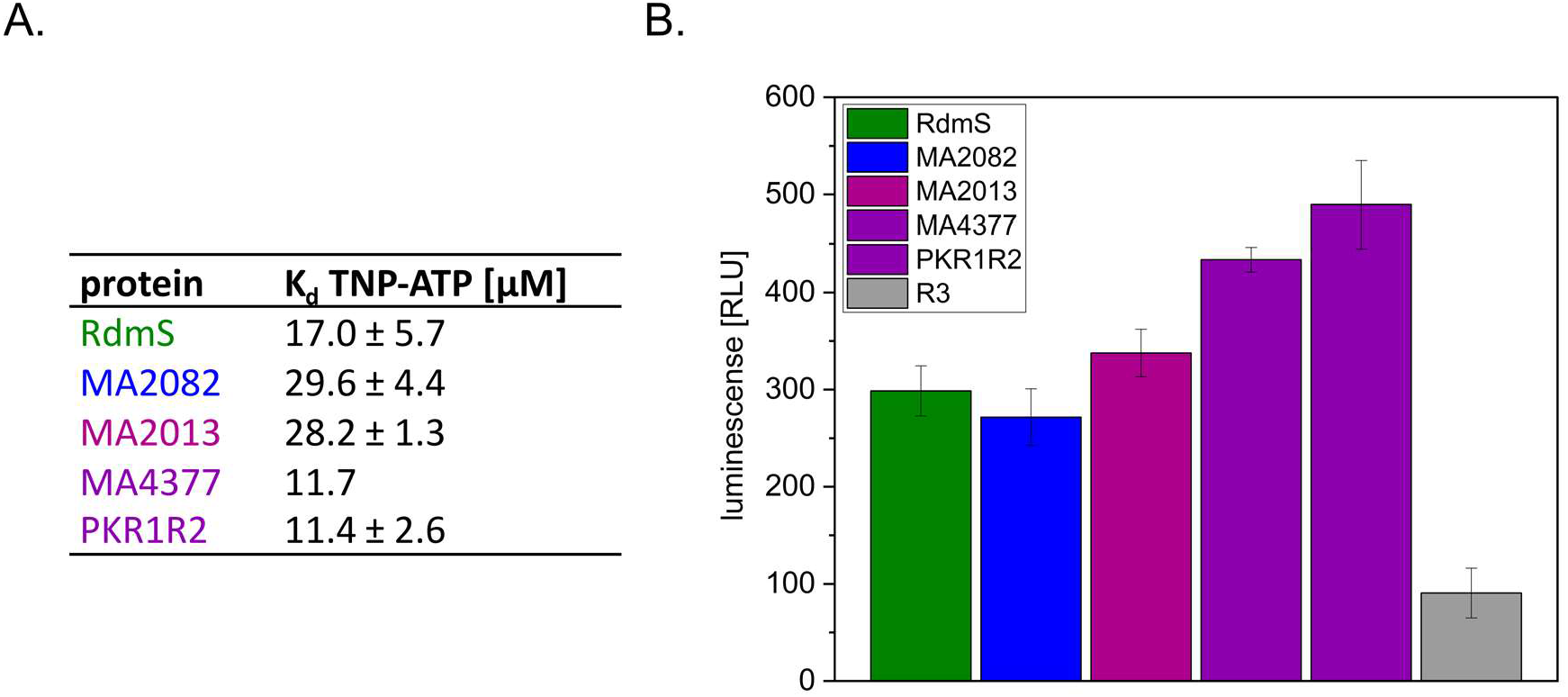
Analysis of ATP binding and hydrolysis. (**A**.) K_d_ values of different putative HKs determined for TNP-ATP. Data were generated using the TNP-ATP assay. Emission spectra were recorded for different protein concentrations (2.5 µM, 5 µM, 7.5 µM) and TNP-ATP concentrations (0 – 50 µM) in kinase buffer (see also Fig. S4). The fluorescence maximum at 541 nm was plotted against the TNP-ATP concentration for every protein concentration. K_d_ values were examined using the SOLVER function of Microsoft Excel. The results were averaged over three independent measurements. **(B.)** ADP-glo assay to examine the hydrolysis of ATP for different HK. R3, the REC-only RR in vicinity to MA4377 was used as negative control since it is a non-ATP binding protein. The mean values and standard deviations of three independent kinase reactions are shown. RLU: relative light unit.

### RdmS (MA_type 2 kinase) is not an autokinase

*In vitro* autokinase activity was tested using recombinant proteins and radiolabelled [γ-^32^P] ATP. Although all four proteins showed ATP binding and hydrolysis, not all of them showed autokinase activity. As previously reported, the RdmS kinase (MA_type 2) lacking the H-box (Figure 1F) did not show autophosphorylation in kinase assays under fully reducing conditions. However, in our previous study, autophosphorylation of a tyrosine residue was reported under oxidising conditions (Fiege & Frankenberg-Dinkel, 2019). Reinvestigation of this observation revealed that the kinase assays under oxidising conditions are only positive when the reaction is terminated with a 4x SDS sample buffer lacking a reducing reagent (i.e. β-mercaptoethanol). Furthermore, this autophosphorylation is dependent on the presence of cysteine residues in the protein and appears to be an unspecific phosphorylation phenomenon which also occurs in non-kinase proteins (Supplement Figure S4A and S4B). To summarise, it must be revised that RdmS is a redox-dependent autokinase but is instead a kinase with an unknown substrate.

### Representatives of MA_type 1 and 3 kinases are true histidine kinases

The cytosolic kinase MA2082, representative of MA_type 3 kinases, and MA2013, the cytosolic representative of MA_type 1B, clearly show time-dependent autokinase activity (Supplement Figure S4C and S4D). As both proteins possess an H-box with a putative phospho-accepting His residue, MA2082 and MA2013 are likely true HKs.

The MA4377 full-length protein (membrane representative of MA_type 1A) (Figure 3A.1) and a variant without the bound REC domains (Figure 3A.2) did not show autokinase activity (Supplement Figure S4E). Different purification methods for the membrane protein (nanodiscs and detergent) had no influence on the autokinase activity, leaving us with the question whether MA4377 is a true HK or not. Therefore, we decided to test truncated cytosolic variants of the protein to rule out any problems associated with the membrane protein purification. A variant only consisting of the PAS and HK domain (designated PK; Figure 3A.5) indeed displayed time-dependent autokinase activity, confirming MA4377 also is a putative HK (Figure 3B). In addition to PK, the recombinant protein versions PKR1R2 and PKR1 (Figure 3A.3 and 3A.4) were incubated with [γ-^32^P] ATP and showed clear autokinase activity (Supplement Figure S4F and S4G). To verify H497 as the conserved histidine residue, the protein variant PK_H497Q_ was generated using site-directed mutagenesis. This protein variant binds ATP but lacks autokinase activity (Figure 3C) proving MA4377 undergoes phosphorylation at this His residue.

**Figure 3:**
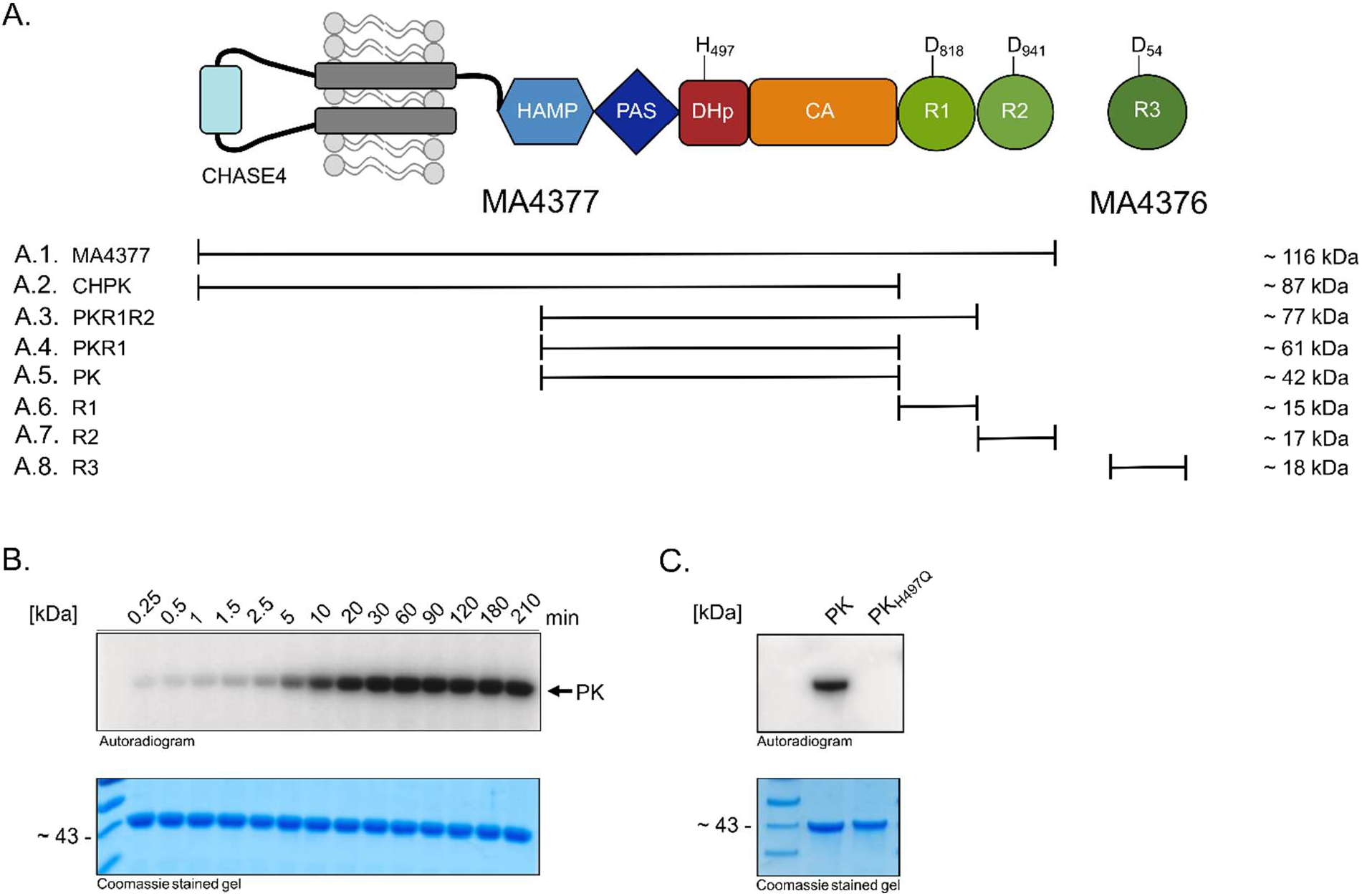
The hybrid histidine kinase MA4377. (**A**.) Schematic representation of the domain structure and overview of truncated versions of the protein. All versions of the protein MA4377 used in this study are shown with their molecular weight. The sites of phosphorylation are highlighted in one letter amino acid code (H497, D818, D940 and D54). (**A.1**.) Full length membrane protein MA4377, (**A.2**.) MA4377 without C-terminal REC domains, (**A.3**.) MA4377 without the CHASE and HAMP domain, (**A.4**.) MA4377 without the CHASE, HAMP and R2 domains, (**A.5**.) MA4377 without CHASE4, HAMP, R1 and R2 domains, (**A.6**.) only the R1 domain of MA4377, (**A.7**.) only the R2 domain of MA4377. (**A.8**.) REC-only RR MA4376 (R3). (**B**.) Autophosphorylation assay with PK. Purified recombinant PK was phosphorylated with [γ-^32^P]-ATP and the reaction was stopped at indicated time points using 4x SDS sample buffer. Samples were separated via SDS-PAGE and subjected to autoradiography (Autoradiogram) or Coomassie staining to visualise the loaded protein (Coomassie stained gel). (**C**.) Identification of the histidine phosphorylation site. Analysis of autophosphorylation activity of the conserved histidine residue variant PK_H497Q_, [γ-^32^P]-ATP radiolabelled samples were separated by SDS-PAGE and the radioactive signals detected by PhosphorImager (Autoradiogram). The same SDS gel was stained with Coomassie (Coomassie stained gel) to visualize the loaded protein.

### MA4377 is part of a phosphorelay in *M. acetivorans*

MA4377 is the most complex kinase selected for our study with a periplasmic CHASE4 domain which is linked to a cytoplasmic HAMP (**H**istidine kinases, **A**denylate cyclases, **M**ethyl accepting proteins and **P**hosphatases) and PAS domain (Figure 3A). The central HK domain is C-terminally fused to two REC domains (R1 and R2). In addition, a REC-only RR (R3) is encoded downstream (MA_4376), overlapping with MA_4377 by 29 nucleotides. The complexity of MA4377 suggests a putative role in a phosphorelay system which has not been studied in Archaea thus far. We therefore decided to explore its function further and to biochemically investigate the phosphorylation cascade.

First, we asked whether the fused REC domains R1 and R2 as well as R3 are involved in a phosphorelay mediated by MA4377. Therefore, the protein variant PK (Figure 3A.5) was autophosphorylated and the transphosphorylation to R1 (using construct PK_H497Q_R1, see Figure 4D) and R2 (using construct PK_H497Q_R1_D818N_R2, see Figure 4D) as well as to R3 (Figure 3A.8) was explored.

**Figure 4:**
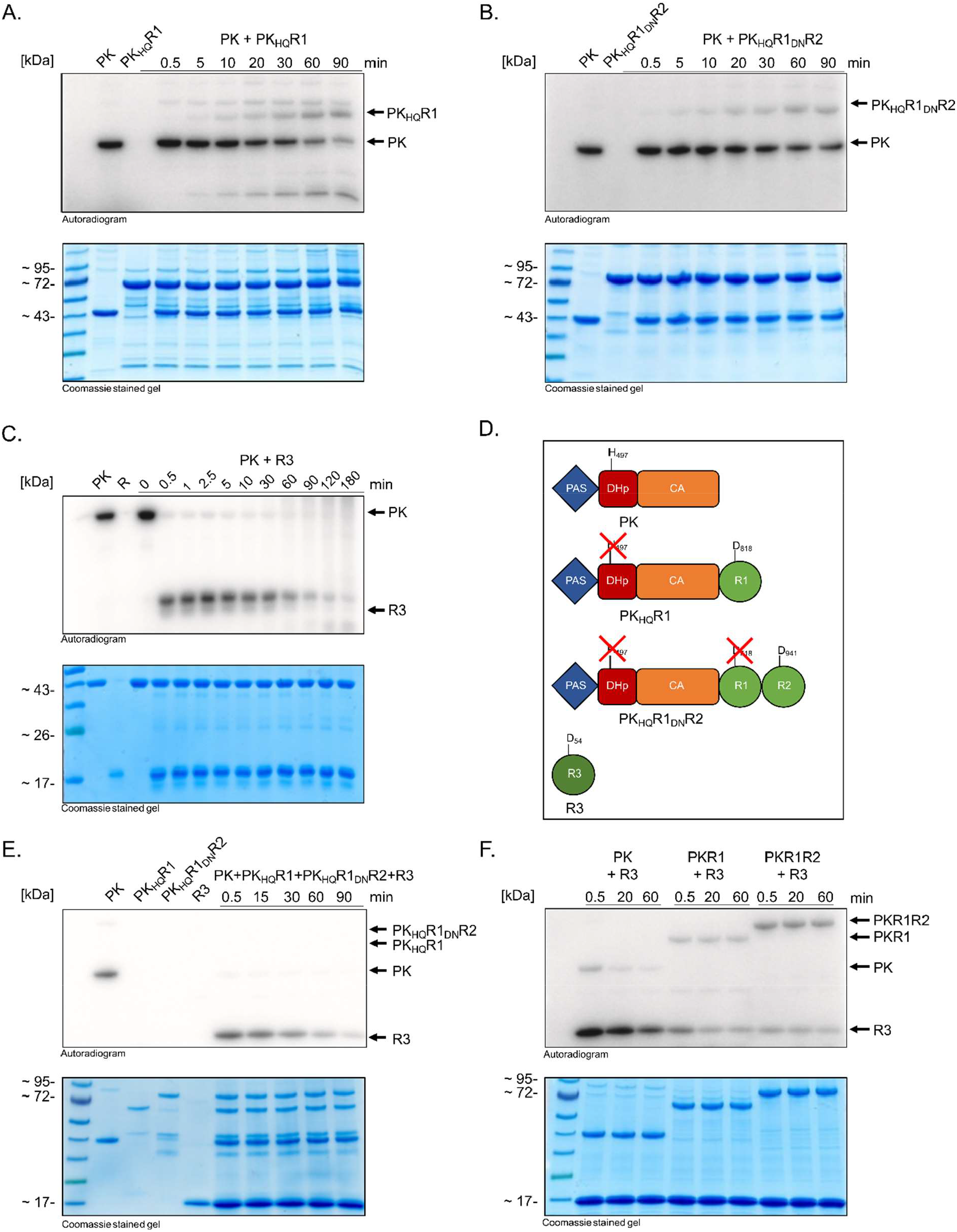
Transphosphorylation assays of PK variants of MA4377. Autoradiogram and corresponding SDS-PAGE (Coomassie stained gel) of the transphosphorylation reaction of different protein variants. Purified recombinant PK (10 µM) was phosphorylated with [γ-^32^P]-ATP, after removing excessive ATP with illustra™ MicroSpin™ G-25 Columns (GE healthcare), the second protein was added in equimolar amount. (**A**.) Transphosphorylation assay with PK and PK_H497Q_R1. (**B**.) Transphosphorylation assay of PK with PK_H497Q_R1R2. (**C**.) Transphosphorylation of PK with R3. (**D**.) schematic representation of applied protein variants. (**E**.) Transphosphorylation Assay with PK and PK_H497Q_R1, PK_H497Q_R1_D818N_R2, R3. PK was autophosphorylated and all constructs were added simultaneously. The H497 and D818 residue were replaced by glutamine and asparagine, respectively via site directed mutagenesis to prevent autophosphorylation of the construct and in case of PK_H497Q_R1_D818N_R2 to additional prevent phosphate transfer to R1. (**F**.) Transphosphorylation assay with different MA4377 variants (PK, PKR1, PKR1R2). The signal that corresponds to the autophosphorylated kinase is indicated by an arrow.

Time-dependent transphosphorylation was observed to both bound REC domains (R1 and R2) and additional to the REC-only R3 (Figure 4A-C, Supplement Figure S5). Immediately after incubation, a fast transfer of the radioactive signal from PK to R3 was observed, while the R3 itself showed no autophosphorylation activity. At first glance, the transphosphorylation of PK to R3 seems to be faster than to the bound REC domains R1 and R2 (Figure 4C). We therefore decided to check whether there is a preference for the transfer. When PK is autophosphorylated and all three constructs (PK_H497Q_R1, PK_H497Q_R1_D818N_R2, R3) were added simultaneously, a signal was only visible at R3 indicating that the kinase preferable transfers phosphate to R3 (Figure 4E). Another experiment with kinase variants with unmodified (i.e. no site directed variants) bound REC domains showed a similar result. PK shows almost complete transphosphorylation to R3. However, if R1 and R1R2 are also present in the MA4377 protein, the signal from the kinase does not disappear completely, suggesting that radiolabelled phosphoryl-group internally went to R1 and or R2 (Figure 4F). Finally, all three REC domains are involved in a phosphorelay of MA4377, but R3 seems to be the preferred target to the kinase.

### Phosphoprofile of MA4377 give hints for cross regulation of HK and RRs in *M. acetivorans*

As REC domains typically display a high degree of structural similarity, the question arose on whether the observed transfer to R1, R2, and R3 is specific or if other RRs are also involved in the phosphorelay. *M. acetivorans* encodes a total of 15 RRs, 11 of them are so called REC-only RRs, only consisting of a REC domain, and four are fused to domains with unknown function. For 13 RRs, transphosphorylation assays were performed to investigate whether they are involved in the phosphorelay mediated by MA4377. The remaining two RR were excluded as they are associated with CheA type HK. The protein variants PK and PKR1R2 were autophosphorylated and, after removing excessive ATP, the different RRs were added. Fast transphosphorylation from phosphorylated PK to MA2445 and R3 was observed within 30 seconds (Figure 5A). After 60 minutes, transphosphorylation to MA2861, MA4671, R1 and R2 was additionally visible (Figure 5B). In a second experiment, the autophosphorylated variant PKR1R2 was used. This experiment verified the transphosphorylation of MA2445 and R3 (Figure 5C). For MA2861 and MA4671 the transfer is ambiguous, as the signals are just slightly visible after 60 minutes (Figure 5D).

**Figure 5:**
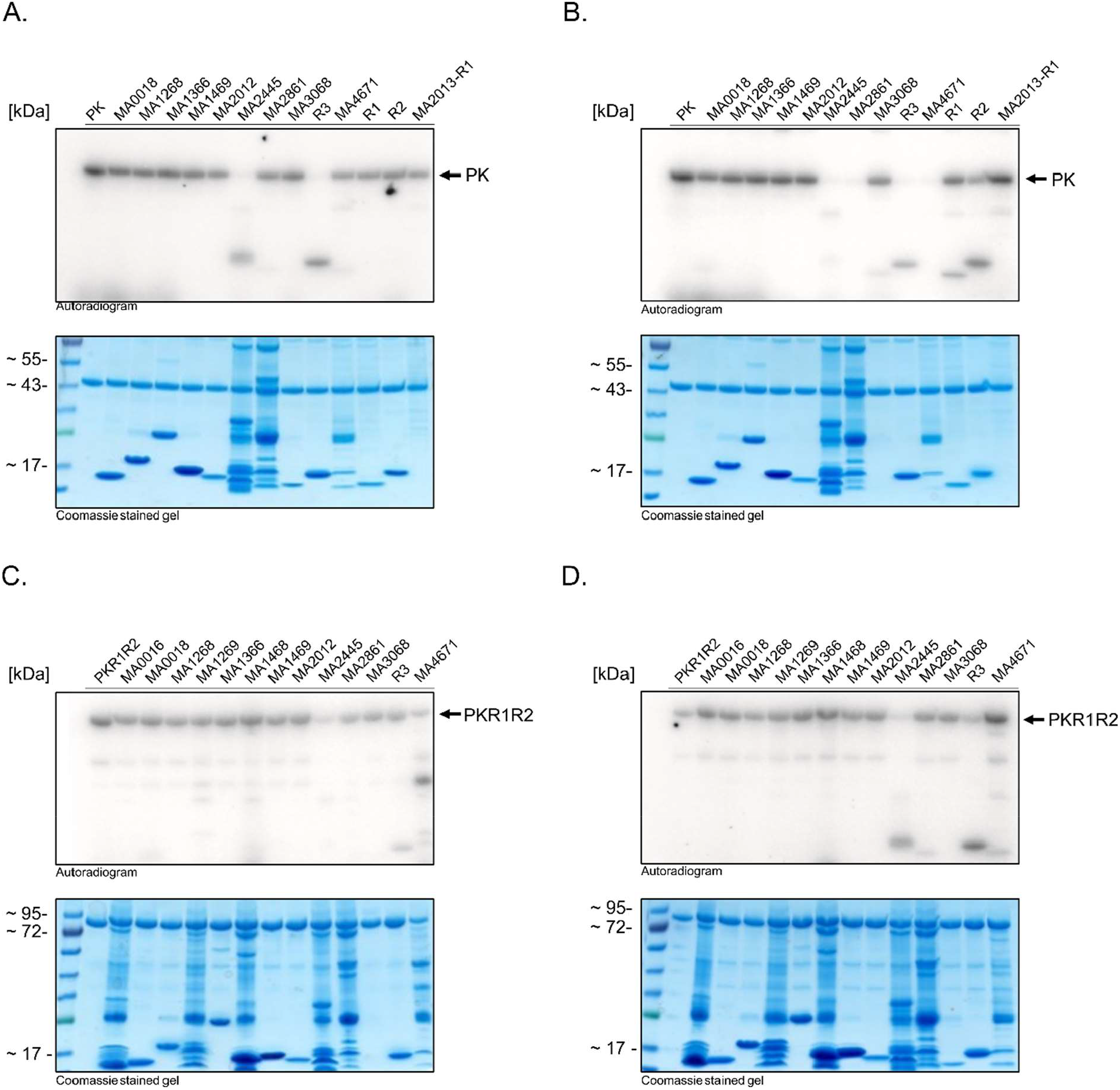
Phosphotransfer profiling of PK and PKR1R2 variants of MA4377. The purified 10 µM of the HKs PK and PKR1R2 were incubated with radioactive labelled [γ-^32^P]-ATP, after removing excessive ATP with illustra™ MicroSpin™ G-25 Columns (GE healthcare), the second protein (RR) was added in equimolar amount and the reaction was stopped after 30 seconds or 60 minutes using 4x SDS sample buffer with β-mercaptoethanol. Samples were separated by SDS-PAGE and the radioactive signals detected by PhosphorImager (Autoradiogram). The same SDS gel was stained with Coomassie (Coomassie stained gel) to visualize the loaded protein (**A**.) Phosphoprofile of PK after 30 seconds. (**B**.) Phosphoprofile after 60 minutes. (**C**.) Phosphoprofile of PKR1R2 after 30 seconds. (**D**.) Phosphoprofile after 60 minutes. The signal that corresponds to the autophosphorylated kinase is indicated by an arrow.

**Figure 6:**
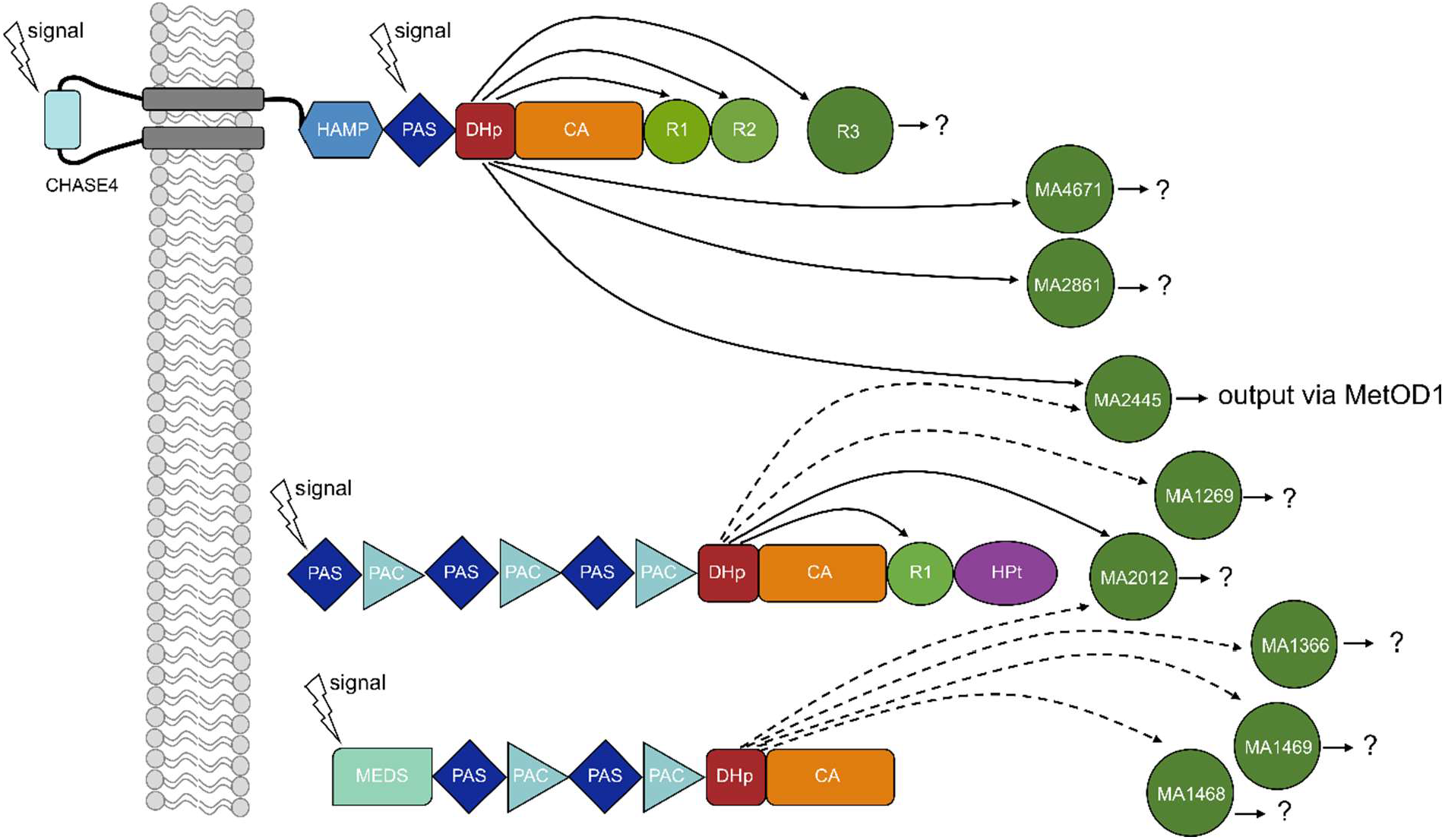
Schematic representation of a putative phosphorelay systems of *M. acetivorans* based on the three putative kinases MA4377, MA2013 and MA2082. Signals can be sensed from the outside and the inside of the membrane. black lines represent verified transfer, dashed lines represent possible transfer.

This experiment showed that besides the bound REC domains R1 and R2 and the nearby REC-only protein R3, also other RRs are able to receive phosphate from MA4377. This suggests a complex phosphorelay system involving MA4377.

### Possible further steps of the phosphorelay of MA4377

In a classical bacterial phosphorelay system, the phosphoryl group is transferred from the His residue of the kinase to an Asp residue in the REC domain. This then targets a His residue of a HPt domain containing protein, finally ending up at an Asp of a RR with output domain. Based on our results for the putative phosphorelay system of MA4377, the next signaling partner should be a HPt domain-containing protein in case the system is similar to those found in bacteria. Therefore, the genome of *M. acetivorans* was screened for HPt domain containing proteins. Overall, we identified three such proteins (MA0014, MA3066 and MA2013). Two of them are CheA-like chemotaxis proteins (MA0014 and MA3066) with a N-terminal HPt domain and are therefore likely not involved in the MA4377 phosphorelay. The third one (MA2013) contains a C-terminal HPt domain. Since the number of HPt domain containing proteins is low, we also searched for regulatory proteins encoded in the genomic vicinity of MA4377. We identified two transcriptional regulators (MsrC and MsrX) with HTH domain and DUF1724 domain. Our future endeavours will therefore target the further downstream signalling experimentally. Preliminary data however, excluded the involvement of the Msr-regulators (Supplement Figure S6).

## Discussion

Sensing and reacting to environmental stimuli via signal transduction systems is a key factor for the survival of microorganisms. While several examples of bacterial and eukaryotic signal transduction systems have been investigated experimentally in the last years, the knowledge of archaeal signal transduction is still limited. Therefore this study gives a first glimpse at the complexity of signal transduction in the methanogenic archaeon *M. acetivorans*.

### HK and RR of *M. acetivorans* show distinct differences to bacterial signal transduction systems

Looking closer at the number and distribution of HK and RR in the genome of *M. acetivorans*, it is evident that kinases outnumber REC proteins by at least three times. In addition, only six of the HK are encoded in an operon with a corresponding RR, resulting in a high number of orphan kinases. This is almost opposite to the situation of TCS in *E. coli*. There, 89 % of the HK are encoded in operons with their corresponding RR (Capra & Laub, 2012). The high number of HK suggests that signal transduction in *M. acetivorans* does not work in strict pairs and that cross-regulation exists (Galperin *et al*., 2018; Krell, 2018). Alternatively, *M. acetivorans* might use alternative but unknown signalling partners for its orphan kinases. Another interesting result from the *in silico* analysis is the number of predicted membrane-bound HK. Most bacterial HK are membrane bound (around 73 %) (Wuichet *et al*., 2010), compared to archaea that exhibit around 38 % membrane-bound HK (Galperin *et al*., 2018), *M. acetivorans* only has about 23 % predicted membrane HK. This observation already gives a hint on the putative signals to be sensed. Membrane-bound kinases are often required to sense signals that cannot easily enter the cell and are usually external signals, while cytosolic signal input domains are used for detecting internal signals or those that can easily pass the membrane (i.e. light, gasses). Despite the differences in membrane bound HK, the overall domain organisation is largely similar to bacterial HK with PAS and GAF domains being highly prevalent (Galperin *et al*., 2018; Koretke *et al*., 2000; Krell, 2018).

Looking closer into the RR of *M. acetivorans*, it is noticable that they differ from bacterial RR. In bacterial signal transduction, the signal receiving REC domain is often linked to DNA binding domains (such as helix-turn helix (HTH) motifs) which play a role in transcriptional regulation (Galperin, 2006; Hoch & Varughese, 2001). Overall, there is only one RR with HTH domain in the genome of *M. acetivorans* (MA1366). On the other hand, a high number of REC-only RRs, without any output domain are present. This is not only the case for the methanogenic archaeon investigated in this study but an overall phenomenon within the Archaea suggesting that other signal transduction mechansims have evolved (Galperin, 2006, 2010). Additional novel output domains occur that are structural and functional undescribed (halobacterial output domain (HalOD) and methanogen output domain (MetOD) (Galperin *et al*., 2018).This leads to the assumption that transcriptional regulation might not be the primary function of archaeal signal transduction systems (Krell, 2018). Besides transcriptional regulation, bacterial RR are known to contribute to ligand binding, protein binding or enzymatic reactions (Galperin, 2006, 2010). REC-only RR can produce an output directly through a conformational change in the REC domain (Koshland, 1977; Stewart *et al*., 2000). A prominent example for an REC-only RR is CheY that works together with CheA and CheB in the chemotaxis signalling in *E. coli*. Phosphorylated CheY binds to the flagella motor protein FliM which controls the direction of flagellar rotation (Muok *et al*., 2020; A. Stock *et al*., 1988). Furthermore, REC-only proteins are also known as phosphate sinks and thereby can serve as buffers in complex phosphorelay systems (Galperin, 2006; Szurmant & Ordal, 2004).

### All new defindes kinase types are true autokinases except MA_type 2

Based on the phylogeny of the H-box containing HK domain, we defined four new types of HK in *M. acetivorans*. Our experimental data support the conclusion that all, except the tested member of MA_type 2 are true autokinases, most likely representing HKs as they all possess a conserved His residue within the H-box (Figure 1F, Supplement Figure S2). MA_type 2 kinases completely lack the H-box region but are able to bind and hydrolyse ATP suggesting that they might phosphorylate another signaling partner, possibly a protein partner similar to what is realised in MAP kinase signaling in Eukarya (Pearson *et al*., 2001). However, more experimental data will be required to proof this hypothesis.

### MA4377 is a hybrid kinase central to a complex phosphorelay system

Within this work we provide experimental evidence for the first phosphorelay system in *M. acetivorans*. The hybrid kinase MA4377 was shown to be a HK and to be involved in intra- and intermolecular transphosphorylation of REC domains with the downstream encoded R3 REC-protein being the preferred phosphate acceptor. However, we also showed that intramolecular transphosphorylation still takes place and R1 and R2 are still being phosphorylated, even if the reaction seems to be slower than to R3. Such preferences are also known from bacterial phosphorelay systems which show diverse options for the preference and involvement of RECs/RR. For instance, the chemotaxis system of *Rhizobium meliloti* (CheA/CheY1/CheY2) is using one of the two REC-only CheY RR as a phosphate sink. But only the contribution of both CheY proteins leads to a functional reaction (Sourjik & Schmitt, 1998). The kinase RodK from *Mycococcus xanthus* and PcrK from *Xanthomonas campestris* are examples that not all present REC domains are actively involved in the phosphorelay (Wang *et al*., 2017; Wegener-Feldbrügge & Søgaard-Andersen, 2009).

The phosphoprofil for MA4377 revealed that additional to R1, R2 and R3 also MA2445, MA2861 and MA4671 might be involved in a phopshorelay of MA4377. MA2861 and MA4671 are also REC-only proteins, while MA2445 is a RR with MetOD1 domain. While MA4671 is encoded in an operon with the HHK MA2256 and therefore constitute a signalling system, MA2445 and MA2861 are stand-alone orphan RR. Overall, these data suggest that MA4377 is involved in a complex phosphorelay system. That this system is not the only one in *M. acetivorans* is supported by our phosphoprofile data for the HK MA2082 and MA2013, which also indicate more complex phosphorelay systems with multiple components (Supplement Figure S7).

Our hypothesis is that MA4377 can sense different input signals and, accordingly, react with the transfer of phosphate to different RR. One way could be a TCS of MA4377 and MA2445 with a direct output of the MetOD1 domain. Another way could involve R1, R2 and R3 with either an unknown HPt-like component and a final output, or a direct reaction through a conformational change. Furthermore, one could think of protein:protein interaction directly influencing the function of the protein partner (i.e. an enzymatic reaction). The high number of RR suggests that those proteins are the key players in archaeal signal transduction and a coordination hub for processing and forwarding signals. Overall, our data only provide a small glimpse into the complexity of signal transduction in *M. acetivorans* and Archaea in general. More experimental data including genetic studies are needed to understand the complexity of archaeal signalling in detail.

## Material & Methods

### Phylogenetic analysis

Protein sequences for the phylogenetic trees were taken from Uniport. Accession numbers of the used sequences are found in Supplement Table S4. The alignment for the phylogenetic tree was created with Clustal Omega (Sievers *et al*., 2011). The maximum-likelihood phylogenetic tree was constructed using PhyML 3.0 (Guindon *et al*., 2010) with the WAG substitution model for amino acids (Whelan & Goldman, 2001) and a bootstrap replication of 100, bootstrap values were calculated using the online tool BOOSTER *(Lemoine et al*., *2018)*. Bootstrap values are displayed as light blue circles in different sizes. To generate the tree, only the HK domain of the HK were considered.

### Bacterial strains, media and growth conditions

Descriptions of each strain used in this study are given in Supplement Table S1. All *E*.*coli* strains used in this study were grown with shaking in LB-medium. For solid media LB-Agar was used. When necessary, media were supplemented with antibiotics at the following concentrations: ampicilin (Amp 100 mg/ml) or chloramphenicol (Cm 34 mg/ml).

### Construction of expression plasmids and site-directed mutagenesis

Expression plasmids were constructed by Gibson Assembly cloning. The corresponding genes were amplified by PCR from genomic DNA of *M. acetivorans* WWM73 that was grown in HS medium supplemented with 125 mM methanol. For the plasmids pASK-IBA3_MA4377PK_H497Q_ and pASK-IBA3_MA4377PK_H497Q_R1 synthetic genes were used.

Amino acid variants were generated by QuikChange™ site-directed mutagenesis with Phusion Polymerase and using mismatched primer pairs and the corresponding plasmid template. Used plasmids are listed in Supplement Table S2 and corresponding oligonucleotides are listed in Supplement Table S3.

### Protein production and purification

To produce the recombinant proteins cultures were grown at 37 °C in 1 L LB-medium to an OD_600_ of ∼0.6 and gently cooled down to 17 °C. Expression of pET21a(+)and pACYC-duet1 vectors were induced with 0.5 mM isopropyl β-thiogalactoside (IPTG), pASK-IBA3 vectors were induced with 200 ng/µl anhydrotetracycline (AHT). Cultures of either promoter system were further incubated for 16 h at 17 °C and harvested for 15 min at 3500 g at 4 °C. The cell pellet was stored at −20 °C or directly used for protein purification.

For protein purification cells were resuspended in lysis buffer (for Strep: 100 mM Tris/HCl, 300 mM NaCl, pH 8; for His: 100 mM Tris/HCl, 300 mM NaCl, 10 mM Imidazol pH 7.5), 2 mL buffer per g of cells. Furthermore, 0.25 mM 4-(2-aminoethyl)benzene-sulfonyl fluoride (AEBSF) and 1 mM 1,4-dithiothreitol (DTT), DNAse and lysozyme was added. For membrane proteins the protease inhibitiors:1 μM Leupeptin, 1 μM E-64, 0,15 μM Aprotinin und 500 μM PMSF and 10 μl DNase (5 mg/ml) were added additionally. The cells were lysed with a cell fluidizer (Microfluidizer) with three times 15000 PSI. Lysed cells were centrifuged (Sorvall Lynx Sorvall Lynx 6000 Centrifuge; Rotor: T29-8×50) for 45 min with 50000 g at 4 °C. For cytosolic proteins, the protein lysate was directly applied onto affinity-chromatography column for purification. For membrane proteins the lysed cells were centrifuged for 15 min with 9000 g at 4 °C. The resulting lysate was centrifuged for 1 h with 80000 g at 4 °C, the pellet was resuspended and solubilized with either 1 % DDM (Dodecyl-β-D-maltosid) (Carl Roth) for 2 h or with 10 % DIBMA (diisobutylene-maleic acid) (Glycon Biochemicals) over night at 4 °C. After the solubilisation the protein solution was centrifuged for 1 h with 70000 g at 4 °C and the resulting lysate was applied to the affinity-chromatography column. For Strep-tagged proteins, Strep-Tactin Sepharose (IBA Lifescience) resin was used and for His-tagged proteins TALON superflow (Cytivia) resin was used. For Strep-tagged membrane proteins the Strep-Tactin XT 4 Flow high-capacity resin (IBA Lifescience) was used. Both resins were used according to the instructions of the manufacturer. Elution fractions were dialysed in kinase buffer (50 mM Tris/HCL, 150 mM NaCl, 5 mM MgCl_2_, 50 mM KCl, 10 % Glycerol, pH 8) and concentrated using Amicon concentrator devices (Merck Millipore). Kinase buffer was supplemented with 0.25 % DDM in case of membrane purification.

### TNP-ATP Assay

To investigate the ATP binding affinity of the putative HK, the ATP derivative 2’,3’-O-trinitrophenyl adenosine 5’-triphosphate (TNP-ATP) was used. TNP-ATP has the advantage over native ATP that it can be analysed using fluorescence spectroscopy, as the fluorescence in aqueous solution is very low, but increases with binding to the nucleotide binding site of proteins (Hiratsuka, 2003). To investigate the binding of TNP-ATP, an increasing concentration of TNP-ATP (0 - 80 µM) was mixed with the putative HKs (2.5 μM, 5 μM and 7.5 μM) in kinase buffer. A fluorescence emission spectrum in the range of 450 - 650 nm was recorded using a fluorescence spectrometer (FP-8300 fluorescence spectrometer, Jasco) and a quartz cuvette (SUPRASIL® cuvette, 3×3 mm, Hellma Analytics). The excitation wavelength was 410 nm and the slit width was 5 nm. The fluorescence of TNP-ATP was determined as a maximum at 541 nm. The measurements were carried out in triplicates and the fluorescence emission values determined corrected by subtracting the buffer emission values and the protein emission values determined under the same conditions. To determine the binding affinity, the recorded emission maxima were plotted against the TNP-ATP concentrations. The K_d_ value was determined using the SOLVER function of Microsoft Excel.

### ADP glo Assay

To investigate whether the putative HKs can hydrolyse ATP the ADP Glo^™^ Kinase Assay (Promega) was performed according to the manufacturer’s instructions.

The kinase reaction was carried out in white 96-well plates (Greiner LUMITRACTM 200) and detection was done using a luminometer (FLUOstar® Omega, BMG Labtech). All measurements were carried out in triplicate and the luminescence values determined were corrected by subtracting the values determined under the same conditions from a blank sample. A standard curve was created to quantify the amount of ADP converted during the kinase reaction.

### Radioactive *in vitro* autophosphorylation and transphosphorylation assays

For autophosphorylation assays 10 µM of the purified HK in kinase buffer was incubated by the addition of 200 μM ATP and 3.3 μM [γ-32P]-ATP (3000 Ci/mmol and 10 mCi/ml; Hartmann Analytic) in a final volume of 12.5 μl to start the reaction. For transphosphorylation assays the kinase was autophosphorylated as described before and excessive ATP was removed using illustra™ MicroSpin™ G-25 Columns (GE healthcare). Afterwards the phosphate receptor protein (10 µM) was added to the reaction mixture. Samples were taken after different timepoints. To stop the reaction 12.5 µl of the reaction mixture was added to 2.5 µl of 4 x SDS-sample buffer (with β-mercaptethanol). Samples were separated by SDS-PAGE gels (percentage depending on the size of the proteins) and imaged using a Typhoon FLA 7000 PhosphorImager (GE Healthcare).

## Abbreviations

ATP: adenosintriphosphat
CHASE4: **C**yclases/**H**istidine kinases **A**ssociated **S**ensory **E**xtracellular
DHp: **D**imerisation and **H**istidine **p**hosphotransfer
GAF: c**G**MP-specific phosphodiesterases, **a**denylyl cyclases and **F**hlA
HAMP: **H**istidine kinases, **A**denylate cyclases, **M**ethyl accepting proteins and **P**hosphatases
HHK: Hybrid histidine kinase
HK: Histidine kinase
HPt: **H**istidine **P**hospho**t**ransfer
MEDS: **ME**thanogen/methylotroph **D**cmR **S**ensory
OCS: One component system
PAC: **P**AS-**A**ssociated **C**-terminal motif
PAS: **P**er-**A**rnt-**S**IM
R/REC: **R**eceiver domain
RR: Response regulator
TCS: Two component system
TNP-ATP: 2’-(3’)-O-(2,4,6-trinitrophenyl)-adenosin 5’-triphosphat

## Author contributions

**Nora FK Georgiev:** Conceptualisation, Formal analysis, Investigation, Methodology, Writing-original draft. **Anne L Andersson:** Investigation, Methodology. **Zoe Ruppe:** Investigation. **Loriana Kattwinkel:** Investigation, Methodology. **Nicole Frankenberg-Dinkel:** Conceptualisation, Funding acquisition, Project administration, Writing-review & editing, Supervision.

## Acknowledgements

This work was in part supported by a grant from the Deutsche Forschungsgemeinschaft to NFD. LK was supported by a PhD fellowship by the Carl Zeiss Foundation.

We would like to thank Christine Förster-Schorr (RPTU Kaiserslautern) for her excellent technical assistance, Eugenio Perez Patallo (RPTU Kaiserslautern) for his help with establishing the membrane purification method and Federica Frascogna (RPTU Kaiserslautern) for critically reading the manuscript. Additional thanks go to Prof. Dr. Kirsten Jung (LMU München) for providing the *E. coli TKR2000* strain and helpful discussion regarding membrane protein purification and Dr. Stefan Jacob (IBWF Mainz) for helpful discussions about signal transduction systems.

