## supplementary material for "Archaeal signalling networks - new insights into the structure and function of histidine kinases and response regulators of the methanogenic archaeon *Methanosarcina acetivorans*"

**Table S1 Prokaryotic strains**

| Strain | Genotype | Reference |
|---|---|---|
| *E. coli* Strains | | |
| *E. coli* DH5α | $F^-$ *endA1 glnV44 thi-1 recA1 relA1 gyrA96 deoR nupG purB20* φ80d*lacZ*ΔM15 Δ(*lacZYA-argF*)U169, *hsd*R17($r_K^-$ $m_K^+$), $\lambda^-$ | (Hanahan, 1983) |
| *E. coli* Bl21 (DE3) | $F^-$ *ompT gal dcm lon hsdS*$_B$($r_B^-$ $m_B^-$) λ(DE3 [*lacI lacUV5-T7p07 ind1 sam7 nin5*]) [*malB*$^+$]$_{K-12}$($\lambda^S$) | (Studier & Moffatt, 1986) |
| *E. coli* Nissle 1917 | serotype O6:K5:H1 | (Grozdanov *et al.*, 2004) |
| *E. coli* C43 (DE3) | $F^-$ *omp*T *hsdS*B($r_B^-$ $m_B^-$) *gal dcm* (DE3) | (Miroux & Walker, 1996) |
| *M. acetivorans* strains | | |
| *M. acetivorans* WWM73 | Δ*hpt*::P*mcrB-tetR*-ϕC31-*int*-attP | (Guss *et al.*, 2008) |

**Table S2: Vectors and plasmids used in this study**

| Plasmid | Characteristic | Reference |
|---|---|---|
| pASK-IBA3 | Expression vector, heterologous overexpression in *E. coli*, Strep-tag II, *tet* promotor, $Amp^R$ | IBA Lifescience |
| pACYCDuet-1 | Expression vector, heterologous overexpression and coexpression in *E. coli*, His-tag and S-tag, T7 promoter, $Cm^R$ | Novogene |
| pET21a(+) | Expression vector, heterologous overexpression in *E. coli*, T7-tag and His-tag, T7 promotor, $Amp^R$ | Novogene |
| pASK-IBA3-MA_4377 | pASK-IBA3 derivate, coding region of MA_4377 from *M. acetivorans* at *Sac*II/*Nco*I with C-terminal Strep-tag II, *tet* promoter, $Amp^R$ | (Sexauer, 2021) |
| pASK-IBA3-MA_4377-CHPK | pASK-IBA3 derivate, coding region of MA_4377 (CHASE-HK domain) from *M. acetivorans* at *Bam*HI/*Nco*I with C-terminal Strep-tag II, *tet* promoter, $Amp^R$ | This study |
| pASK-IBA3-MA_4377-PKR1R2 | pASK-IBA3 derivate, coding region of truncated cytosolic MA_4377 (HK-R2 domain) from *M. acetivorans* with C-terminal Strep-tag II, *tet* promoter, $Amp^R$ | (Sexauer, 2021) |
| pASK-IBA3-MA_4377-$PK_{H497Q}R1_{D818N}R2$ | pASK-IBA3-MA_4377-PKR1R2 derivate with amino acid exchange: His497 to Gln and Asp818 to Asn, obtained by site-directed mutagenesis | This study |
| pASK-IBA3-MA_4377-PKR1 | pASK-IBA3 derivate, coding region of truncated cytosolic MA_4377 (HK-R1 domain) from *M. acetivorans* at *Bam*HI/*Nco*I with C-terminal Strep-tag II, *tet* promoter, $Amp^R$ | This study |
| pASK-IBA3-MA4_377-$PK_{H497Q}R1$ | pASK-IBA3-MA4_377-$PK_{H497Q}R1$ derivate with amino acid exchange: His497 to Gln, synthetic gene | This study |
| pASK-IBA3-MA4_377-PK | pASK-IBA3 derivate, coding region of truncated cytosolic MA_4377 (only HK domain) from *M. acetivorans* with C-terminal Strep-tag II, *tet* promoter, $Amp^R$ | (Sexauer, 2021) |
| pASK-IBA3-MA4_377-$PK_{H497Q}$ | pASK-IBA3-MA_4377-PK derivate with amino acid exchange: His497 to Gln, Synthetic gene | This study |
| pASK-IBA3-MA_4377-R1 | pASK-IBA3 derivate, coding region of truncated cytosolic MA_4377 (only R1 domain) from *M. acetivorans* at *Xba*I/*Nco*I with C-terminal Strep-tag II, *tet* promoter, $Amp^R$ | This study |
| pASK-IBA3-MA_4377-R2 | pASK-IBA3 derivate, coding region of truncated cytosolic MA_4377 (only R2 domain) from *M. acetivorans* at *Xba*I/*Nco*I with C-terminal Strep-tag II, *tet* promoter, $Amp^R$ | This study |

| | | |
|---|---|---|
| pASK-IBA3-MA_0863(rdmS)$_{O216K}$ | pASK-IBA3 derivate, coding region of MA_0863 (rdmS) with amino acid exchange (Pyl216 to Lys) from *M. acetivorans* at *Sac*II/*Pst*I with C-terminal Strep-tag II, *tet* promoter, Amp$^R$ | (Kwiatkowski, 2013) |
| pASK-IBA3-MA_2082 | pASK-IBA3 derivate, coding region of MA_2082 from *M. acetivorans* at *Bam*HI/*Nco*I with C-terminal Strep-tag II, *tet* promoter, Amp$^R$ | This study |
| pASK-IBA3-MA_2013 | pASK-IBA3 derivate, coding region of MA_2013 from *M. acetivorans* at *Bam*HI/*Nco*I with C-terminal Strep-tag II, *tet* promoter, Amp$^R$ | This study |
| pACYC-duet-MA_2013-PK | pACYCDuet-1 derivate, coding region of truncated MA_2013 (without R1 and HPt) from *M. acetivorans* at *Bam*HI/*Nde*I with N-terminal His-tag, T7 promoter, Cm$^R$ | This study |
| pASK-IBA3-MA_2013-R1 | pASK-IBA3 derivate, coding region of truncated MA_2013 (only R1) from *M. acetivorans* with C-terminal Strep-tag II, *tet* promoter, Amp$^R$ | This study |
| pASK-IBA3-MA_2013-HPT | pASK-IBA3 derivate, coding region of truncated MA_2013 (only HPt) from *M. acetivorans* with C-terminal Strep-tag II, *tet* promoter, Amp$^R$ | This study |
| pACYC-duet1-MA_0016 | pACYCDuet-1 derivate, coding region of MA_0016 from *M. acetivorans* at *Bam*HI/*Nde*I with N-terminal His-tag, T7 promoter, Cm$^R$ | This study |
| pACYC-duet1-MA_0018 | pACYCDuet-1 derivate, coding region of MA_0018 from *M. acetivorans* at *Bam*HI/*Nde*I with N-terminal His-tag, T7 promoter, Cm$^R$ | This study |
| pACYC-duet1-MA_1268 | pACYCDuet-1 derivate, coding region of MA_1268 from *M. acetivorans* at *Bam*HI/*Nde*I with N-terminal His-tag, T7 promoter, Cm$^R$ | This study |
| pACYC-duet1-MA_1269 | pACYCDuet-1 derivate, coding region of MA_1269 from *M. acetivorans* at *Bam*HI/*Nde*I with N-terminal His-tag, T7 promoter, Cm$^R$ | This study |
| pACYC-duet1-MA_1366 | pACYCDuet-1 derivate, coding region of MA_1366 from *M. acetivorans* at *Bam*HI/*Nde*I with N-terminal His-tag, T7 promoter, Cm$^R$ | This study |
| pACYC-duet1-MA_1468 | pACYCDuet-1 derivate, coding region of MA_1468 from *M. acetivorans* at *Bam*HI/*Nde*I with N-terminal His-tag, T7 promoter, Cm$^R$ | This study |
| pACYC-duet1-MA_1469 | pACYCDuet-1 derivate, coding region of MA_1469 from *M. acetivorans* at *Bam*HI/*Nde*I with N-terminal His-tag, T7 promoter, Cm$^R$ | This study |
| pACYC-duet1-MA_2445 | pACYCDuet-1 derivate, coding region of MA_2445 from *M. acetivorans* at *Bam*HI/*Nde*I with N-terminal His-tag, T7 promoter, Cm$^R$ | This study |
| pACYC-duet1-MA_2012 | pACYCDuet-1 derivate, coding region of MA_2012 from *M. acetivorans* at *Bam*HI/*Nde*I with N-terminal His-tag, T7 promoter, Cm$^R$ | This study |

| | | |
|---|---|---|
| pACYC-duet1-MA_2861 | pACYCDuet-1 derivate, coding region of MA_2861 from *M. acetivorans* at *Bam*HI/*Nde*I with N-terminal His-tag, T7 promoter, $Cm^R$ | This study |
| pACYC-duet1-MA_3068 | pACYCDuet-1 derivate, coding region of MA_3068 from *M. acetivorans* at *Bam*HI/*Nde*I with N-terminal His-tag, T7 promoter, $Cm^R$ | This study |
| pACYC-duet1-MA_4376 (R3) | pACYCDuet-1 derivate, coding region of MA_4376 from *M. acetivorans* at *Sac*I/*Sal*I with N-terminal His-tag, T7 promoter, $Cm^R$ | (Sexauer, 2021) |
| pACYC-duet1-MA_4671 | pACYCDuet-1 derivate, coding region of MA_4671 from *M. acetivorans* at *Bam*HI/*Nde*I with N-terminal His-tag, T7 promoter, $Cm^R$ | This study |
| pET21a(+)-MA_4375 (*msrX*) | pET21a (+) derivate, coding region of MA_4375 from *M. acetivorans* with C-terminal His-tag, T7 promoter, $Amp^R$ | (Sexauer, 2021) |

**Table S3: Oligonucleotids used in this study**

| Oligonucleotids | Sequence (5' → 3') |
|---|---|
| **Oligonucleotids for sequencing** | |
| pASK-IBA3-seq-fwd | CGCAGTAGCGGTAAACG |
| pASK-IBA3-seq-rev | GAGTTATTTACCACTCCCT |
| pACYC-duet1-seq-fwd | TCTCCCTTATGCGACTCCTG |
| pACYC-duet1-seq-rev | GGGTTATGCTAGTTATTGCTCAGC |
| **Oligonucleotids for expression plasmids** | |
| pASK-IBA3-MA_4377-fwd | GCTATCCGCGGCATGAATGTGAGTAGAAAAATTCT |
| pASK-IBA3-MA_4377-rev | GCATACCATGGCCTCTTCGACAATAAGAATTTCTTTC |
| pASK-IBA3-MA_4377-CHPK-fwd | ATTCGAGCTCGGTACCCGGGATATGAATGTGAGTAGAAAAATTC |
| pASK-IBA3-MA_4377-CHPK-rev | GTGGCTCCAAGCGCTGAGACTCTGAGCTTCCCTATTTTC |
| pASK-IBA3-MA_4377-PKR1R2-fwd | GCTATCCGCGGCAGGTTGAATTCCGATAAGGTTAA |
| pASK-IBA3-MA_4377-PKR1R2-rev | GCATACCATGGCCTCTTCGACAATAAGAATTTCTTTC |
| pASK-IBA3-MA_4377-PKR1-fwd | ATTCGAGCTCGGTACCCGGGCTAGGTTGAATTCCGATAAG |
| pASK-IBA3-MA_4377-PKR1-rev | GTGGCTCCAAGCGCTGAGACGGAAACTGAACTTACCTG |
| pASK-IBA3-MA_4377-PK-fwd | GCTATCCGCGGCAGGTTGAATTCCGATAAGGTTAA |
| pASK-IBA3-MA_4377-PK-rev | GCGCCATGGCCTGTGAGGGGAATTG |
| pASK-IBA3-MA_4377-PK für QC | ATTCGAGCTCGGTACCCGGGCTATGAGGTTGAATTCCG |
| pASK-IBA3-MA_4377-PK für QC | GTGGCTCCAAGCGCTGAGACGCTTTGTGAGGGGAATTG |
| pASK-IBA-MA_4377-R1-fwd | ATGAATAGTTCGACAAAAATTAGTCTTGTGCTTGTAGTC |
| pASK-IBA3-MA_4377-R1-rev | GTGGCTCCAAGCGCTGAGACTGAAACTGAACTTACCTG |
| pASK-IBA3-MA_4377-R2-fwd | ATGAATAGTTCGACAAAAATTGGTAAGTTCAGTTTCGAC |
| pASK-IBA3-MA_4377-R2-rev | GTGGCTCCAAGCGCTGAGACTTCTTCGACAATAAGAATTTCTTTC |
| pASK-IBA3-MA_0863(rdmS)$_{O216K}$-fwd | GGACCGCGGAAAAAAGTTCGATATAAATC |

| | |
|---|---|
| pASK-IBA3-MA_0863(rdmS)$_{O216K}$-rev | GGACTGCAGTTATTTTTCGAACTCCGGGTGGCTCCAAGACTCTATTTGATTCATG |
| pASK-IBA3-MA_2082-fwd | ATTCGAGCTCGGTACCCGGGATATGGAAGGAAAACTGAGAAATTCTGGTATTGAG |
| pASK-IBA3-MA_2082-rev | GTGGCTCCAAGCGCTGAGACTCTCGACGCCCACAGGGA |
| pASK-IBA3-MA_2013-fwd | ATTCGAGCTCGGTACCCGGGAAGTGTGCGGTTTAAAAAAAC |
| pASK-IBA3-MA_2013-rev | GTGGCTCCAAGCGCTGAGACGACAGATCTGTTTTTCCAG |
| pACYC-duet1-MA_2013-PK-fwd | ACCATCATCACCACAGCCAGATGTGCGGTTTAAAAAAAC |
| pACYC-duet1-MA_2013-PK-rev | TATCCAATTGAGATCTGCCACTAGCATACTAACAGGCTTTC |
| pASK-IBA3-MA_2013-R1-fwd | ATTCGAGCTCGGTACCCGGGCGATGGTGAAAAGAGGGG |
| pASK-IBA3-MA_2013-R1-rev | GTGGCTCCAAGCGCTGAGACCTATACTTTCCAGGTCCAG |
| pACYC-duet1-MA_0016-fwd | ACCATCATCACCACAGCCAGATGGCAAGAGTAATGATC |
| pACYC-duet1-MA_0016-rev | TATCCAATTGAGATCTGCCATTAATTTGGTTCAGCAATTTTTTTAATTAC |
| pACYC-duet1-MA_0018-fwd | ACCATCATCACCACAGCCAGATGCCTGAAATCCTGATC |
| pACYC-duet1-MA_0018-rev | TATCCAATTGAGATCTGCCATCAACCAGGAGAATTCGTTG |
| pACYC-duet1-MA_1268-fwd | ACCATCATCACCACAGCCAGATGAAAACAAAGGTAGCTG |
| pACYC-duet1-MA_1268-rev | TATCCAATTGAGATCTGCCATTATTTTGACGGTAGTTTCAC |
| pACYC-duet1-MA_1269-fwd | ACCATCATCACCACAGCCAGATGGAAACATGGACGGCATTTAAAC |
| pACYC-duet1-MA_1269-rev | TATCCAATTGAGATCTGCCATTAAGTAAGATTTACGACTTCAAGCC |
| pACYC-duet1-MA_1366-fwd | ACCATCATCACCACAGCCAGATGCATAACATCAATCTTGAC |
| pACYC-duet1-MA_1366-rev | TATCCAATTGAGATCTGCCATCAAATCCGACTACGTCTC |
| pACYC-duet1-MA_1468-fwd | ACCATCATCACCACAGCCAGATGGACAAAGCAAAGATTC |
| pACYC-duet1-MA_1468-rev | TATCCAATTGAGATCTGCCATTATTCTTCAGTTACTTTACCC |
| pACYC-duet1-MA_1469-fwd | ACCATCATCACCACAGCCAGATGAAAAAAGCAAAAATTCTGGTTGTTG |

| | |
|---|---|
| pACYC-duet1-MA_1469-rev | TATCCAATTGAGATCTGCCATCATTCGTCGCTGTCGGAG |
| pACYC-duet1-MA_2012-fwd | ACCATCATCACCACAGCCAGATGAAAGTTCTGATTGCC |
| pACYC-duet1-MA_2012-rev | TATCCAATTGAGATCTGCCATTAAAAGCCCAGCTTTTTTC |
| pACYC-duet1-MA_2445-fwd | ACCATCATCACCACAGCCAGATGTTGAATGAGAAAAACAAAC |
| pACYC-duet1-MA_2445-rev | TATCCAATTGAGATCTGCCATTATGCTGCTTTTATGATAAATTC |
| pACYC-duet1-MA_2861-fwd | ACCATCATCACCACAGCCAGATGAAAGTACTGCTTGTAG |
| pACYC-duet1-MA_2861-rev | TATCCAATTGAGATCTGCCATTAACCTGAACCATTTTTAAC |
| pACYC-duet1-MA_3068-fwd | ACCATCATCACCACAGCCAGATGGCAAAAGTCTTGATTG |
| pACYC-duet1-MA_3068-rev | TATCCAATTGAGATCTGCCATTATGCACCGATTACTTTC |
| pACYC-duet1-MA_4376(R3)-fwd | GCGTCGAGCTCGAAAGAAATTCTTATTGTCG |
| pACYC-duet1-MA_4376(R3)-rev | CCCTTAGTCGACTTCTCCCAGATATTTATCG |
| pACYC-duet1-MA_4671-fwd | ACCATCATCACCACAGCCAGATGAAAAGTATACTTGTAGTTGAG |
| pACYC-duet1-MA_4671-rev | TATCCAATTGAGATCTGCCATCACAATACCTGGTAACC |
| pET21a(+)-MA_4375(msrX)-fwd | CTTTAAGAAGGAGATATACAATGAAAATATCACTTATTGACCTG |
| pET21a(+)-MA_4375(msrX)-rev | AGTGGTGGTGGTGGTGGTGCTTTATTTTGTCCAGCCGC |
| **Oligonucleotids for site directed mutagenesis** | |
| MA_4377-H497Q-fwd | GAGTTCCTTGCCAATATGAGCCAGGAGCTCCGGACGCCTCTG |
| MA_4377-H497Q-rev | CAGAGGCGTCCGGAGCTCCTGGCTCATATTGGCAAGGAACTC |
| MA_4377-D818N-fwd | CCTGATGTTATCACTCTGAATGTTTTACTTCCCGATACTAGTGGC |
| MA_4377-D818N-rev | GCCACTAGTATCGGGAAGTAAAACATTCAGAGTGATAACATCAGG |
| MA_4377-D941N-fwd | CAGCAGCCGGATATCCTCATCCTCAACCTGCTGATGCCCGAGATAAGT |
| MA_4377-D941N-rev | ACTTATCTCGGGCATCAGCAGGTTGAGGATGAGGATATCCGGCTGCTG |

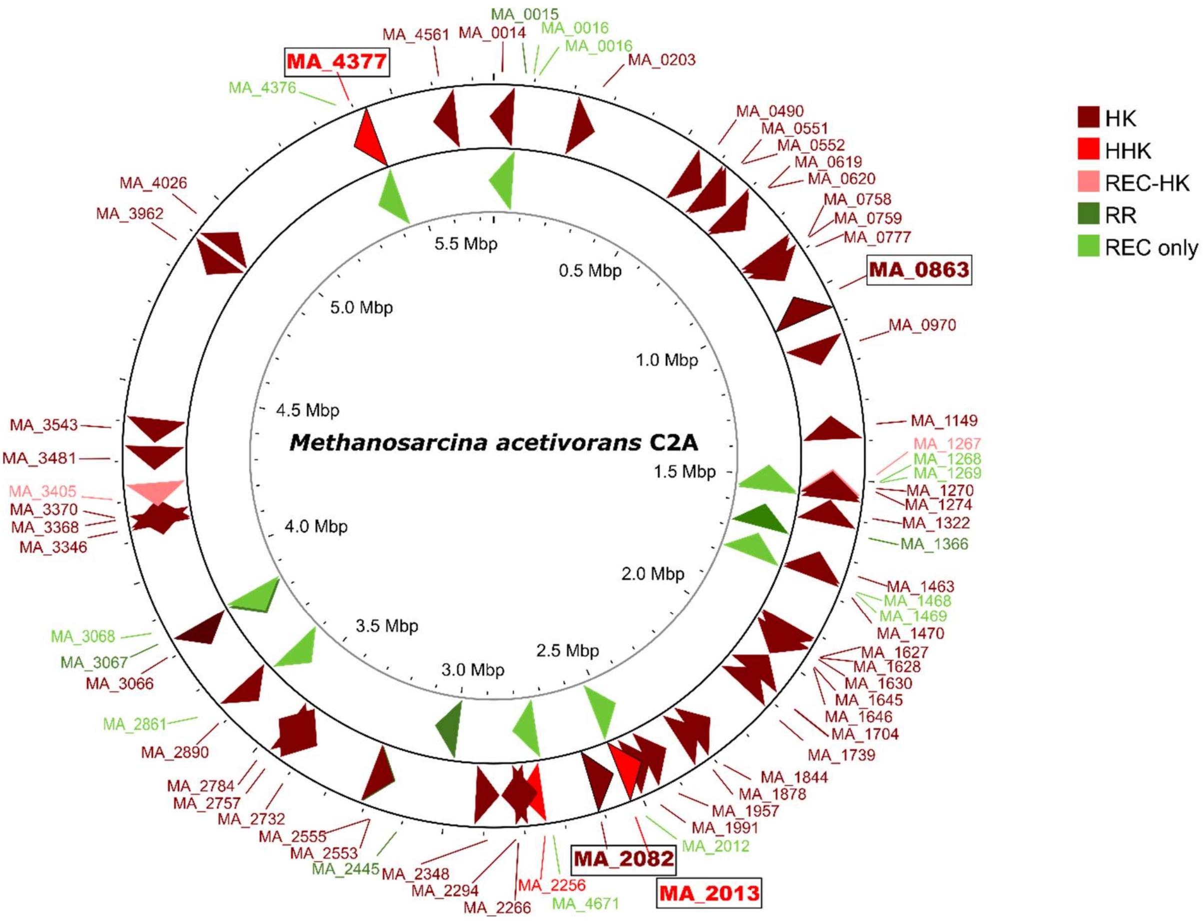


**Figure S1: Distribution of HK and RR in the genome of *M. acetivoans*.** The whole genome of *M. acetivorans* was displayed with PROKSEE regarding the distribution of putative signal transduction components. The different subforms of HK and RR are displayed with different colours. HK = histidine kinase, HHK = hybrid histidine kinase, REC-HK = histidine kinases with N-terminal REC domain, RR = response regulator, REC only = Rec-only response regulator.

```
MA_type 1A   MA3405   NM SH-------ELKTP LNSIIG----F----SDLLKEEIAGPLNEKQSRYVQFISSSGKN  45
             MA2294   SM SH-------ELRTP LNSIIG----F----SDMLLTQNFGPLNKKQLRYVNNISVSGNH  45
             MA3962   NM SH-------ELRTP LNSVIG----F----SDLLLEGAFGPLNTKQSKYVNNILISGKN  45
             MA1149   NM SH-------ELRTP LNAVIG----F----SDLLLSETAGPLNEKQKRYTENISKSGSH  45
             MA2555   NM SH-------ELRTP LNSIIG----F----SDLLYEKIYGDLNEKQLKAVGNISRSGKH  45
             MA1739   NI SH-------ELRTP LNSIIG----F----SDLLCEQIFGELNEKQLRYAGNISKSGKH  45
             MA3368   NM SH-------ELRTP LNSIIG----F----SDMLYEQAYGELNRKQLRAIGNISSSGKH  45
             MA2553   NM SH-------ELRTP LNSIIG----F----SDLLYEKVYGELNLKQTKAVGNISNSGKH  45
             MA4377   NM SH-------ELRTP LNSIIG----F----SDILIERVFGELNEKQLKYVNNISGSGKH  45
             MA2348   NM SH-------ELRTP LNAVIG----F----ADILNEEGFGPLNKKQKRFVGNISTSGKH  45

no type      MA0777   -- -------------- --------------------------------------------   0

MA_type 1B   MA1957   TV SH-------ELKTP LNSIIG----F----SDLLLEDSSGKLSERQARYINNISISGKH  45
             MA2256   TM SH-------ELRTP LTAIIG----F----SELMLGEATGEFDELNRKFLGHISNSGKH  45
             MA2013   NM SH-------EIRTP MNAVLG----M----LEMLLETS---LTDEQREYLQLAHASAES  42

MA_type 1C   MA1270   VS SH-------DLQEP LRMIAS----Y----LQLLQRRYQGELDERADKYIYFAVDGASR  45

MA_type 2    RdmS     -- -------------- ---------------------------EDQQQKAIDTVINSSER  17
             MsmS     EF VE-------EMMFP EK--AE----Y----GEIMDYETLYAIDSQQQKAVNTFIHYSEK  43

MA_type 3    MA0203   EI HH-------RIKNN LQ--------VISSLLDLQAEKFNSREDIKDSEVLEAFRESQDR  45
             MA0490   EI HH-------RIKNN LQ--------VISSLLSLEAEKFSDE------KMLESFRESQNR  39
             MA0551   EI HH-------RIKNN LQ--------VISSLLDLQAEKFQN------KEVLEAFRESQNR  39
             MA0552   EI HH-------RIKNN LQ--------VISSLLDLQAEKFNNREDIKDSEILEAFRESQDR  45
             MA0619   EI HH-------RIKNN LQ--------VISSLLSLEAEKFSDE------RTLEAFRESQNR  39
             MA0620   EI HH-------RIKNN LQ--------VISSLLSLQAEKFEDR------EVIEAFRESQNR  39
             MA0758   EI HH-------RIKNN LQ--------VISSLLDLQAEKFRD------KDVLEAFRESQSR  39
             MA0759   EI HH-------RIKNN LQ--------VISSLLDLQAEKFRD------KEVLEAFRESQSR  39
             MA0970   EI HH-------RIKNN LQ--------VISSLLDLQAEKFRGKKNIEDSKILEAFKESQDR  45
             MA1274   EI HH-------RIKNN LQ--------VISSLLDLQAEKFRSREHVEDSEVLNAFKESQER  45
             MA1322   EI HH-------RIKNN LQ--------VISSLLDLQAEKFGNKKYIMNSEVMDAFRESQDR  45
             MA1470   EI HH-------RIKNN LQ--------VISSLLELQAEKFDNL------EVLEAFRESQNR  39
             MA1627   EI HH-------RIKNN LQ--------VISSLLDLQADKFDNP------KVIEAFRESQNR  39
             MA1628   EI HH-------RIKNN LQ--------IISSLLDLQAEMFKGRSNIRDSEVLKAFAVSMDR  45
             MA1630   EI HH-------RIKNN LQ--------IISSLLDLQAEKFED------KNVTEAFREGQNR  39
             MA1645   EI HH-------RIKNN LQ--------VISSLLDLQAEQFKNRENIKDSEVLEAFRESQDR  45
             MA1646   EL HH-------RIKNN LQ--------VISSLLDLQADLFKGKKTITDSEVLKAFNESIDR  45
             MA1704   EI NH-------RIKNN LQ--------VISSLLSFEAEKSTDP------EILEAFRETQNR  39
             MA1844   EI HH-------RIKNN LQ--------VISSLLELQADKFKDR------EVIEAFRESQNR  39
             MA1878   EI HH-------RIKNN LQ--------VISSLLDLQIDIFSNREICKTPEVIEAFRESQNR  45
             MA1991   EI HH-------RIKNN LQ--------VISSLLSLQAEKFRD------QEVLEAFRESQDR  39
             MA2082   EI HH-------RIKNN LQ--------VISSLLDLQAEKFRD------KEVLEAFRESQNR  39
             MA2266   EI HH-------RIKNN LQ--------VISSLLSLQAEYFSDP------KVKESFKDSQNR  39
             MA2732   EI HH-------RIKNN LQ--------VISSLLDLECDSLLSG-TPDHKKIAEAFRESHNR  44
             MA2757   EI HH-------RIKNN LQ--------VISSLLDLQAEQFKNRECIKNSEVLEAFRESQAR  45
             MA2784   EI HH-------RIKNN LQ--------VISSLLDLQAEKFKDREDIKDSEVLEAFRESQDR  45
             MA3346   EI HH-------RIKNN LQ--------VISSLLDLQAENFSKHEVCKTPKVVEAFKESQDR  45
             MA3370   EI HH-------RIKNN LQ--------VISSLLDLQAGKFNNKEHIRDSEVLEAFKESQDR  45
             MA3481   EI HH-------RIKNN LQ--------VISSMLSLQAEKFSDE------ETLEAFRESQNR  39
             MA3543   EI HH-------RIKNN LQ--------VISSLLDLQAEKFNKREGIKDSEVMEAFRESQDR  45
             MA4026   EI HH-------RIKNN LQ--------VISSLLDLQAEKFSHREAVPTLEILEAFKESQNR  45
             MA1267   EI HH-------RIKNN LQ--------VISSLLDLQAEKFED------PTIRQAFRESQNR  39
             MA1463   EI NH-------RIKNN LQ--------IVSSLLDLQAEQFSDK------KVIEAFRESENR  39
             MA2890   EI HH-------RIKNN LQ--------IVSSLLSLQADKFKDK------DVIEAFRESENR  39

MA_CheA      MA0014   -- -------------- ---------RISTEQLDKL-MNLVGELVINRSRVKELTGESKSK  34
             MA3066   GS SHHFSESAASAKTP LETQRQETIRVRTSNLDNI-MNLVGELVINKGRLLQISQEYNLP  59
```

**Figure S2: Partial amino acid sequence alignment of HisKA domain of all putative HK of *M. acetivorans*.** Acession numbers of the employed sequences are listed in Table S4, for the alignment only the amino acid sequences of the HisKA domain were taken. The Alignment was constructed using CUSTAL Omega and modified using PowerPoint. Conserved His residue is highlighted in bold with gray backgound. H-box region groups into four distinct groups (Ma_type 1 A/B/C, Ma_type 2, Ma_type 3 and MA_CheA). MA_type 1 is similar to the Type I HK of bacteria. As the amino acid arrangement differs within the group, the subgroups A, B and C were created. MA_type 2 kinases are characterized by the absence of an H-box and MA_type 3 kinases cannot be grouped with a known type of bacterial kinases. MA_CheA are highly similar to bacterial CheA proteins.

```
MA0016        --------MARVMIVDDAEFMRMVIRDILLKHGHE-VVAEVGDGEEAIQTYL-------E  44
MA0018        --------MPEILIVEDNLLNLVIEADLLKSCGY--DPKKAKNGFEALEVLS------KV  44
MA1268        MKTKVAAKPIEILLVEDSEGDVGLIEEVFEEAKIRNNLHIVEDGEEAILFLRGEKQFSGI  60
MA1269        METWTAFKPVDILLVEDNKGDVGLIEEVFESSKVRNKLYVVEDGEEAVHFLLREGKFSDV  60
MA1366        ----------KILIMGNGNNVHNNLQKVLEAENY--NVVSASDNFSAIETV-------NE  41
MA1468        ------MDKAKILVVEDQNIVALNLRNRLKNMGYI-VPTIAISGEEAIRKTE-------L  46
MA1469        ------MKKAKILVVEDQNIVALNIRNKLKNLGYT-VPGTASTGEEAIRKAE-------L  46
MA2012        ----------KVLIAEDEPISNLWLKNTLTRWGY--EAISTRDGYEAWEVLN------ES  42
MA2013-R1     ----------NVLFAEDHPINQKLILGLLEKKGH--KLTLVTSGKDALDALS------RR  42
MA2445        ----------KVLIVDDKKENVELMEAYLAVEPY--DVITAYGGKEAFQKV-------KE  41
MA2861        ---------MKVLLVDDDPVFLELSKTFLEVFHDI-NSDTVESARQALEKLD-------E  43
MA3068        --------MAKVLIVDDTAFMRKLLKNILFGAGFD-IAGEAENGKQAVEMYK-------G  44
MA4376        --------MKEILIVEDNPMNMELILDLLEFYGH--RVTEAEDGIKALERLA------EK  44
MA4377-R1     ----------LVLVVDDDINSNELISVVLREAGY--STASLHNGKDVLEVA-------KK  41
MA4377-R2     ----------KVLIIDDDENAVELLSSMIESEGF--EIVKAYSGQAGLDKLF------SE  42
MA4671        ------MEMKSILVVEDSPVILELISFFLTSSGY--ESRETGDGFDALKIAE------EN  46
                        ::.  :             :

MA0016        VKPDLVLMDIIMP-DMDGKEALQKLLLID-----PDAKVVMCSSLGQQALITESMKIGAM  98
MA0018        KVD-LVLMDMELP-KMHGLELLQRIKCNP---ETQGIRVVAVTGHCDPESEQEFLKAGCH  99
MA1268        SRPDIILLDLNLP-KKDGREVLEEIKEDD---DLKNIPVVVLTTSKAEEDVLKSYNLHAN  116
MA1269        PRPGIILLDLNLP-KKDGREVLEEIKEDD---DLKNIPVVVLTTSRAEEDILESYKLHAN  116
MA1366        EKPDLVLLDTVYL-ETDGFEICRQLKDSP---RYWWIPIMMLSERNKTEDGIKAFDSGAD  97
MA1468        TTPDLVLMDIMLKGDMDGIEAARIIKSRF------SAPVIYLTACTDIGILERAKLTEPA  100
MA1469        TNADLVLMDIMLKGDMDGIEAAREIKARL------KIPVLYLTAYTDDETLERAKMTEPA  100
MA2012        DFPRVVILDWEMP-KMKGIEVCEKIKKDP---RLSSIYIILITGRDLTEDMEAGFKAGAD  98
MA2013-R1     DFD-AVLMDIQMP-GMDGLEATRRIRDPSSGVRRHNIPIIAFTARALKEDREKCFEAGMN  100
MA2445        EKPDIILLDVMMP-EVNGYEVCKILKGNP---ETQFIPVLMLTALSELEDRIRGIEVGAD  97
MA2861        LSYDVVVSDYDMP-YMDGISFLKTIRDKR-----INIPFILFTGVGKEEIKSQAIENGVD  97
MA3068        LKPDVVTMDVVMP-EMTGIDALKQIKALD-----KDAKIVMCTAIGQENIVKTAIKLGAR  98
MA4376        KFD-IILLDMQLP-KMDGLEVLDRIKKNP---ATADIPVIAVTAHAMKGSEEHFIEMGCV  99
MA4377-R1     LKPDVITLDVLLP-DTSGWNVLKQLKSDL---DTTSIPVLIISVTDNNE---LGVALGAT  94
MA4377-R2     QQPDILIIDLLMP-EISGFEIISRLRDGE---QTKDIPLIVCTAGEFTEKNIEKLNGELK  98
MA4671        RFD-LILLDKQLP-GFDGLEVLKKIKKIF---EIRKTSVIALMAHAMQGDEDRFLKAGCN  101
                  :  *         * .    :              .:

MA0016        GF-----IIKPFEPDGMLDVIKKIAEPN--------------  121
MA0018        AV-----LSKPINFDLFGAQVKEFLTATNSPG-----------  126
MA1268        AY-----VTKPVDFDQFIRVIKSIEDFWLEVVKLPSK------  148
MA1269        AY-----VTKPVDFDQFIKVIKSIENFWLEVVNLT--------  146
MA1366        DY-----ITMPFNPLELKARVGMIL------------------  117
MA1468        GY-----ISKPFKEKDLVSNIEVALQKNKL-GKVTEE------  131
MA1469        GY-----ISKPFKEEDLHSNIEMALHKHRT-EKKEIENSDSDE  137
MA2012        DY-----LKKPFDNRKLKTKLDTAR------------------  118
MA2013-R1     YY-----ISKPLKKEKLLNILEDIR------------------  120
MA2445        DF-----LTKPINRLELKTRVKSLL------------------  117
MA2861        SL-----IQKRGDPKAQYSELSKRIWQIVKNGSG---------  126
MA3068        GY-----IIKPFQAPKVIEEIKKVIGA----------------  120
MA4376        DY-----ISKPIDIHRFRSLIDKYLGE----------------  121
MA4377-R1     YS-----FTKPVRRVELLDSLREIT------------------  114
MA4377-R2     GHLISIMKKGTFGRKELINRIKQLA------------------  123
MA4671        GY-----ISKPIDIDRFKLILDTCTGGYQVL------------  127
                                  :
```

**Figure S3: Amino acid sequence alignment of REC domain of all putative RR of *M. acetivorans*.** Acession numbers of the employed sequences are listed in Table S4, for the alignment only the amino acid sequences of the REC domain were taken. The Alignment was constructed using CUSTAL Omega and modified using PowerPoint. All proteins share a conserved Asp residue, highlighted in bold with gray background.

**Table S4: Sequences of 53 putative HK and 15 RR of *M. acetivorans*.** The protein name corresponds to the old locus tag. The new locus tag and the acession number of the protein in the uniprot database is displayed in the table. For the Alignment (Figure S2) and the phylogenetic tree (Figure 1A) the aminoacid sequence of the HisKA domain was employed. For the alignment (Figure S3) the amino acid sequence of the REC domain was employed.

| Protein | New locus tag | Organism | Accession number | database |
|---|---|---|---|---|
| MA0014 | MA_RS00065 | *M. acetivorans* | Q8TUQ1 | UniProt |
| MA0203 | MA_RS01090 | *M. acetivorans* | Q8TU70 | UniProt |
| MA0490 | MA_RS02555 | *M. acetivorans* | Q8TTE7 | UniProt |
| MA0551 | MA_RS25000 | *M. acetivorans* | Q8TT86 | UniProt |
| MA0552 | MA_RS02905 | *M. acetivorans* | Q8TT85 | UniProt |
| MA0619 | MA_RS03260 | *M. acetivorans* | Q8TT21 | UniProt |
| MA0620 | MA_RS03265 | *M. acetivorans* | Q8TT20 | UniProt |
| MA0758 | MA_RS03960 | *M. acetivorans* | Q8TSN7 | UniProt |
| MA0759 | MA_RS03965 | *M. acetivorans* | Q8TSN6 | UniProt |
| MA0777 | MA_RS04070 | *M. acetivorans* | Q8TSL9 | UniProt |
| MA0863 (RdmS) | MA_RS04495 | *M. acetivorans* | Q8TSD5 | UniProt |
| MA0970 | MA_RS24390 | *M. acetivorans* | Q8TS36 | UniProt |
| MA1149 | MA_RS05985 | *M. acetivorans* | Q8TRM6 | UniProt |
| MA1267 | MA_RS06580 | *M. acetivorans* | Q8TRB3 | UniProt |
| MA1270 | MA_RS06595 | *M. acetivorans* | Q8TRB0 | UniProt |
| MA1274 | MA_RS06615 | *M. acetivorans* | Q8TRA6 | UniProt |
| MA1322 | MA_RS06850 | *M. acetivorans* | Q8TR62 | UniProt |
| MA1463 | MA_RS24430 | *M. acetivorans* | Q8TQS7 | UniProt |
| MA1470 | MA_RS07625 | *M. acetivorans* | Q8TQS0 | UniProt |
| MA1627 | MA_RS08450 | *M. acetivorans* | Q8TQC1 | UniProt |
| MA1628 | MA_RS08455 | *M. acetivorans* | Q8TQC0 | UniProt |
| MA1630 | MA_RS08465 | *M. acetivorans* | Q8TQB8 | UniProt |
| MA1645 | MA_RS08530 | *M. acetivorans* | Q8TQA5 | UniProt |
| MA1646 | MA_RS08535 | *M. acetivorans* | Q8TQA4 | UniProt |
| MA1704 | MA_RS08850 | *M. acetivorans* | Q8TQ48 | UniProt |
| MA1739 | MA_RS09030 | *M. acetivorans* | Q8TQ13 | UniProt |
| MA1844 | MA_RS24495 | *M. acetivorans* | Q8TPR2 | UniProt |
| MA1878 | MA_RS09795 | *M. acetivorans* | Q8TPM8 | UniProt |
| MA1957 | MA_RS10200 | *M. acetivorans* | Q8TPF6 | UniProt |
| MA1991 | MA_RS24525 | *M. acetivorans* | Q8TPC2 | UniProt |
| MA2013 | MA_RS10480 | *M. acetivorans* | Q8TPA1 | UniProt |
| MA2082 | MA_RS10815 | *M. acetivorans* | Q8TP40 | UniProt |
| MA2256 | MA_RS11710 | *M. acetivorans* | Q8TNM8 | UniProt |
| MA2266 | MA_RS11765 | *M. acetivorans* | Q8TNL9 | UniProt |
| MA2294 | MA_RS11915 | *M. acetivorans* | Q8TNJ1 | UniProt |
| MA2348 | MA_RS12170 | *M. acetivorans* | Q8TNE0 | UniProt |
| MA2553 | MA_RS13275 | *M. acetivorans* | Q8TMU8 | UniProt |
| MA2555 | MA_RS13285 | *M. acetivorans* | Q8TMU6 | UniProt |
| MA2732 | MA_RS14295 | *M. acetivorans* | Q8TMC7 | UniProt |
| MA2757 | MA_RS28770 | *M. acetivorans* | Q8TMA7 | UniProt |
| MA2784 | MA_RS28790 | *M. acetivorans* | Q8TM82 | UniProt |
| MA2890 | MA_RS15160 | *M. acetivorans* | Q8TLY2 | UniProt |

| | | | | |
|---|---|---|---|---|
| MA3066 | MA_RS16035 | *M. acetivorans* | Q8TLH0 | UniProt |
| MA3346 | MA_RS17465 | *M. acetivorans* | Q8TKQ3 | UniProt |
| MA3368 | MA_RS25835 | *M. acetivorans* | Q8TKN3 | UniProt |
| MA3370 | MA_RS24765 | *M. acetivorans* | Q8TKN1 | UniProt |
| MA3405 | MA_RS17780 | *M. acetivorans* | Q8TKK0 | UniProt |
| MA3481 | MA_RS18195 | *M. acetivorans* | Q8TKC7 | UniProt |
| MA3543 | MA_RS24775 | *M. acetivorans* | Q8TK73 | UniProt |
| MA3962 | MA_RS20675 | *M. acetivorans* | Q8TJ26 | UniProt |
| MA4026 | MA_RS21010 | *M. acetivorans* | Q8TIW4 | UniProt |
| MA4377 | MA_RS22890 | *M. acetivorans* | Q8THY1 | UniProt |
| MA4561 (MsmS) | MA_RS23780 | *M. acetivorans* | Q8THF6 | UniProt |
| MA_0016 | MA_RS00075 | *M. acetivorans* | Q8TUP9 | UniProt |
| MA_0018 | MA_RS00090 | *M. acetivorans* | Q8TUP7 | UniProt |
| MA_1268 | MA_RS06585 | *M. acetivorans* | Q8TRB2 | UniProt |
| MA_1269 | MA_RS06590 | *M. acetivorans* | Q8TRB1 | UniProt |
| MA_1366 | MA_RS07090 | *M. acetivorans* | Q8TR18 | UniProt |
| MA_1468 | MA_RS07615 | *M. acetivorans* | Q8TQS2 | UniProt |
| MA_1469 | MA_RS07620 | *M. acetivorans* | Q8TQS1 | UniProt |
| MA_2012 | MA_RS10475 | *M. acetivorans* | Q8TPA2 | UniProt |
| MA_2861 | MA_RS15020 | *M. acetivorans* | Q8TM09 | UniProt |
| MA_3068 | MA_RS16045 | *M. acetivorans* | Q8TLG8 | UniProt |
| MA_4376 | MA_RS22885 | *M. acetivorans* | Q8THY2 | UniProt |
| MA_4671 | MA_RS11705 | *M. acetivorans* | Q8TNM9 | UniProt |
| MA_2445 | MA_RS12700 | *M. acetivorans* | Q8TN48 | UniProt |
| MA_0015 | MA_RS00070 | *M. acetivorans* | Q8TUQ0 | UniProt |
| MA_3057 | MA_RS15990 | *M. acetivorans* | Q8TLH9 | UniProt |

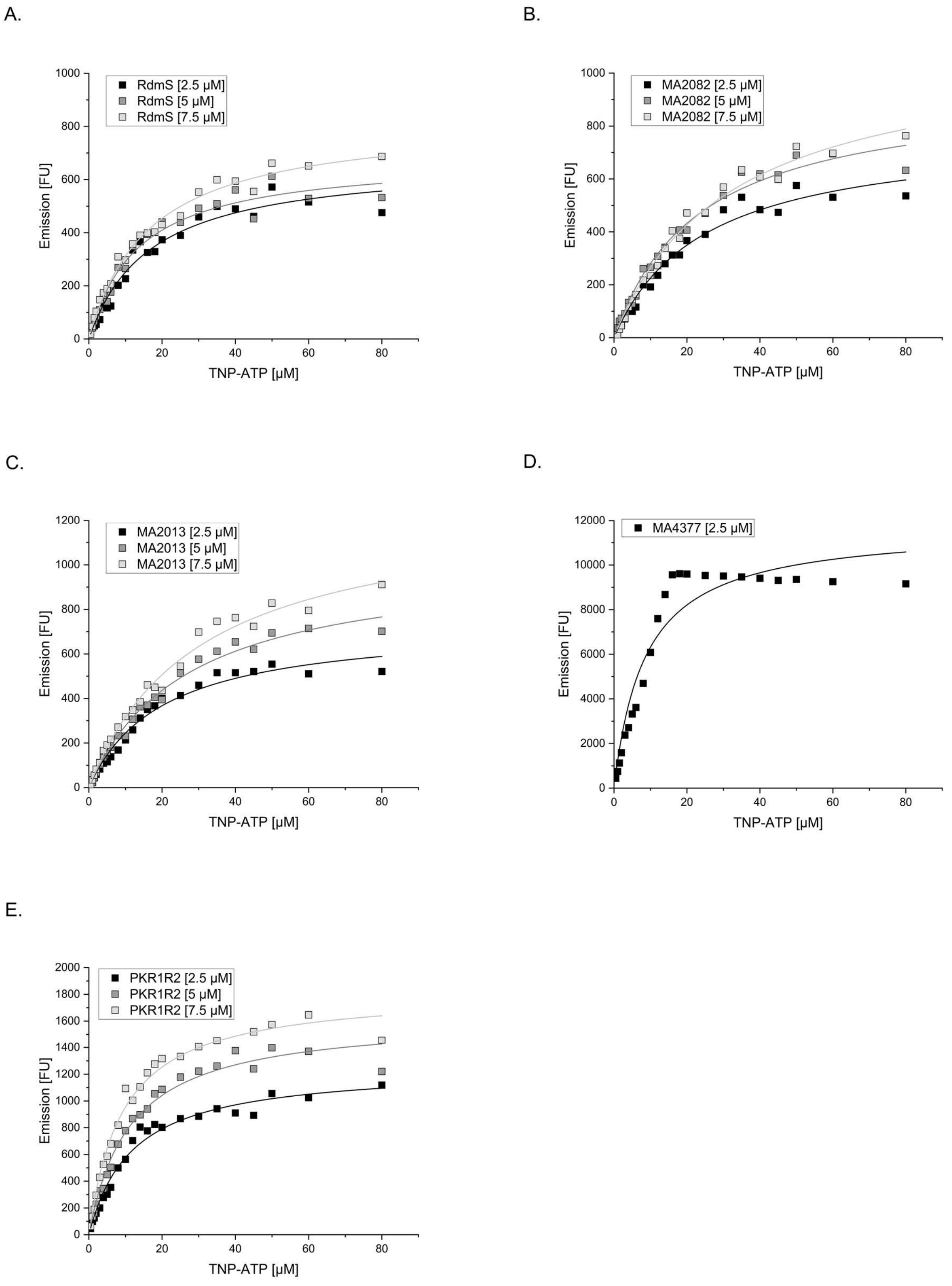


**Figure S4: TNP-ATP association kinetics of putative HKs.** Constant protein concentration (2.5 µM, 5 µM or 7.5 µM) was mixed with variable TNP-ATP concentration (0 – 80 µM) in kinase buffer. A fluorescence emission spectrum in the range of 450 - 650 nm was recorded using a fluorescence spectrometer (FP-8300 fluorescence spectrometer, Jasco) and a quartz cuvette (SUPRASIL® cuvette, 3x3 mm, Hellma Analytics). The excitation wavelength was 410 nm, and the slit width was 5 nm. The fluorescence of TNP-ATP was determined as a maximum at 541 nm. The Kd value was determined using the SOLVER function of Microsoft Excel. (**A.**) TNP-ATP association kinetics of RdmS. (**B.**) TNP-ATP association kinetics of MA2082. (**C.**) TNP-ATP association kinetics of MA2013. (**D.**) TNP-ATP association kinetics of MA4377. (**E.**) TNP-ATP association kinetics of PKR1R2.

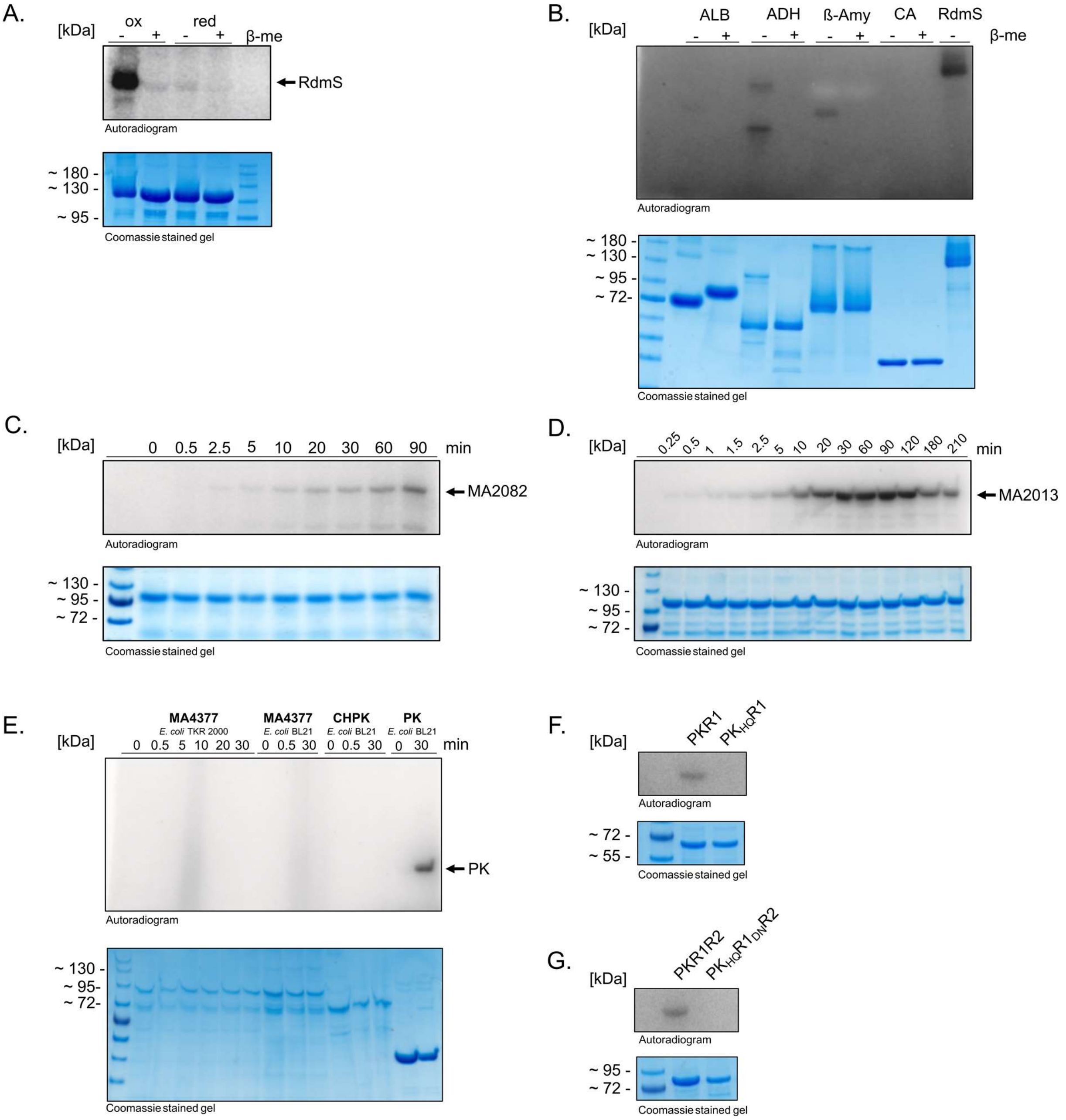


**Figure S5: Autophosphorylation assay of different putative HK.** (**A.**) Autophosphorylation assay of RdmS under oxidizing (ox) and reducing (red) conditions. 10 µM RdmS was incubated with [$\gamma$-$^{32}$P]-ATP and the reaction was stopped using 4x SDS sample buffer with (+) or without (-) β-mercaptoethanol (β-me). (**B.**) Autophosphorylation assay of standard proteins and RdmS under oxidizing conditions. 10 µM of the standard proteins albumin (ALB), alcohol dehydrogenase (ADH), β-amylase (β-Amy), carbonic anhydrase (CA) and RdmS were incubated with [$\gamma$-$^{32}$P]-ATP and the reaction was stopped using 4x SDS sample buffer with (+) or without (-) β-mercaptoethanol (β-me). (**C.**) Autophosphorylation Assay of MA2082. 10 µM MA2082 was incubated with [$\gamma$-$^{32}$P]-ATP and the reaction was stopped using 4x SDS sample buffer with β-mercaptoethanol. (**D.**) Autophosphorylation Assay of MA2013. 10 µM MA2013 was incubated with [$\gamma$-$^{32}$P]-ATP and the reaction was stopped using 4x SDS sample buffer with β-mercaptoethanol. (**E.**) Autophosphorylation Assay of MA4377 and truncated protein variants. MA4377 produced in different *E. coli* strains and truncated variants (CHPK and PK) were incubated with [$\gamma$-$^{32}$P]-ATP and the reaction was stopped using 4x SDS sample buffer with β-mercaptoethanol. (**F.**) Autophosphorylation assay of the MA4377 variants PKR1 and $PK_{H497Q}R1$. 10 µM protein was incubated with [$\gamma$-$^{32}$P]-ATP and the reaction was stopped using 4x SDS sample buffer with β-mercaptoethanol. (**G.**) Autophosphorylation assay of the MA4377 variants PKR1R2 and $PK_{H497Q}R1_{D818N}R2$. 10 µM protein was incubated with [$\gamma$-$^{32}$P]-ATP and the reaction was stopped using 4x SDS sample buffer with β-mercaptoethanol. The samples of all assays were separated by SDS-PAGE and the radioactive signals detected by PhosphorImager (Autoradiogram). The same SDS gel was stained with Coomassie (Coomassie stained gel) to visualize the loaded protein.

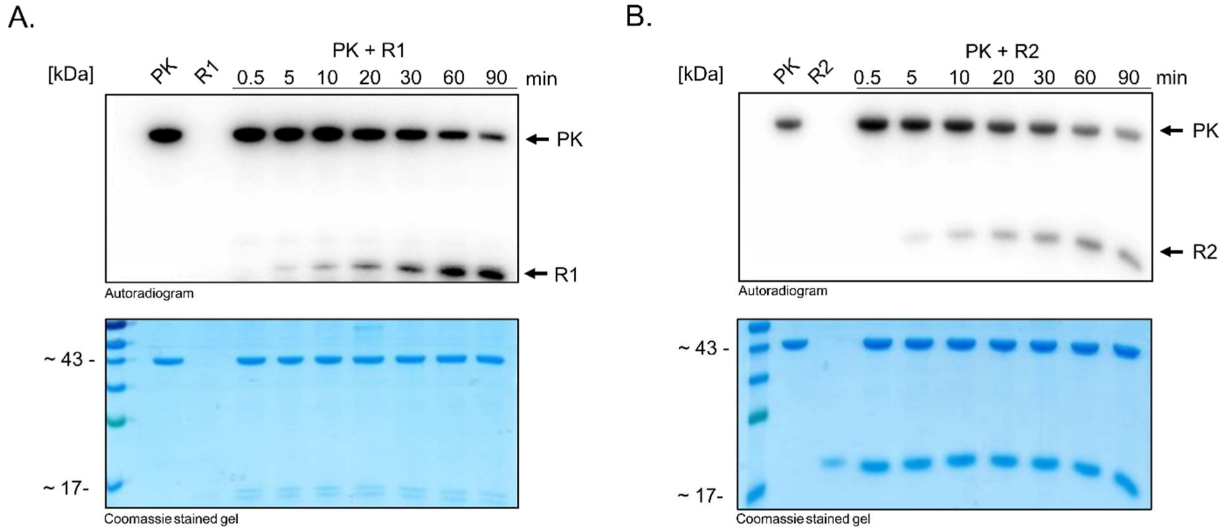


**Figure S6: Transphosphorylation assays of the PK variant of MA4377.** Autoradiogram and corresponding SDS-PAGE (Coomassie stained gel) of the transphosphorylation reaction of different protein variants. Purified recombinant PK (10 µM) was phosphorylated with [γ-$^{32}$P]-ATP, after removing excessive ATP with illustra™ MicroSpin™ G-25 Columns (GE healthcare), the second protein was added in equimolar amount. (**A.**) Transphosphorylation assay with PK and R1. (**B.**) Transphosphorylation assay of PK with R2.

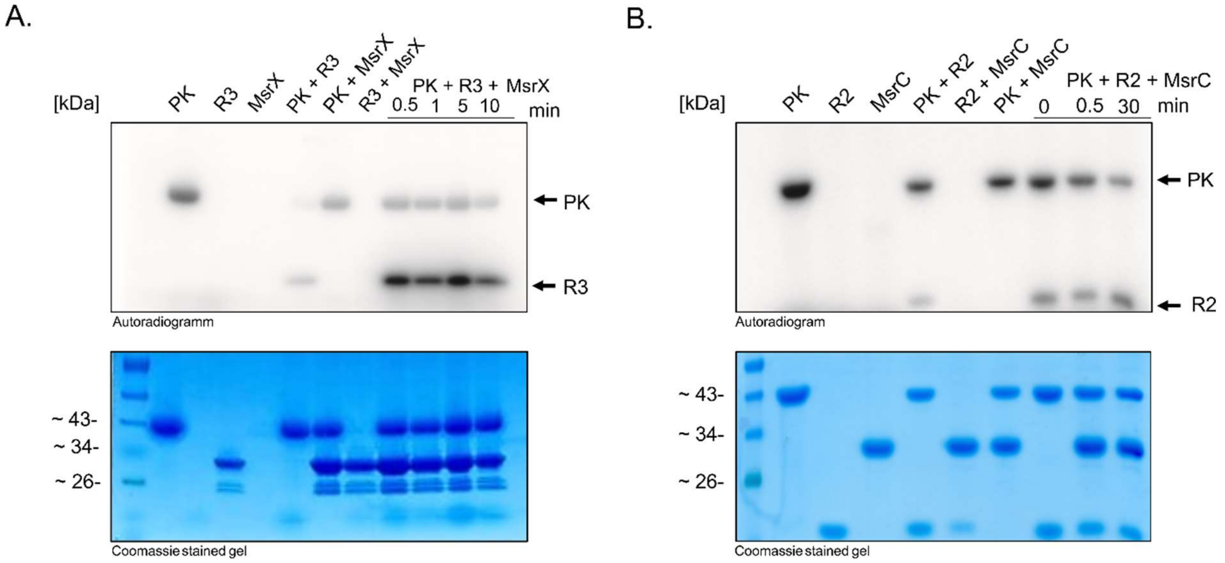


**Figure S7: Putative further components of the phosphorelay of MA4377.** (**A.**) Phosphorylation assay to test whether MsrX is receiving a phosphate from PK or R2. **T**ranscriptional regulator MsrX is not involved in a phosphorelay. (**B.**) Phosphorylation assay to test whether MsrC is receiving phosphate from PK or R3. Transcriptional regulator MsrC is not involved in a phosphorelay.

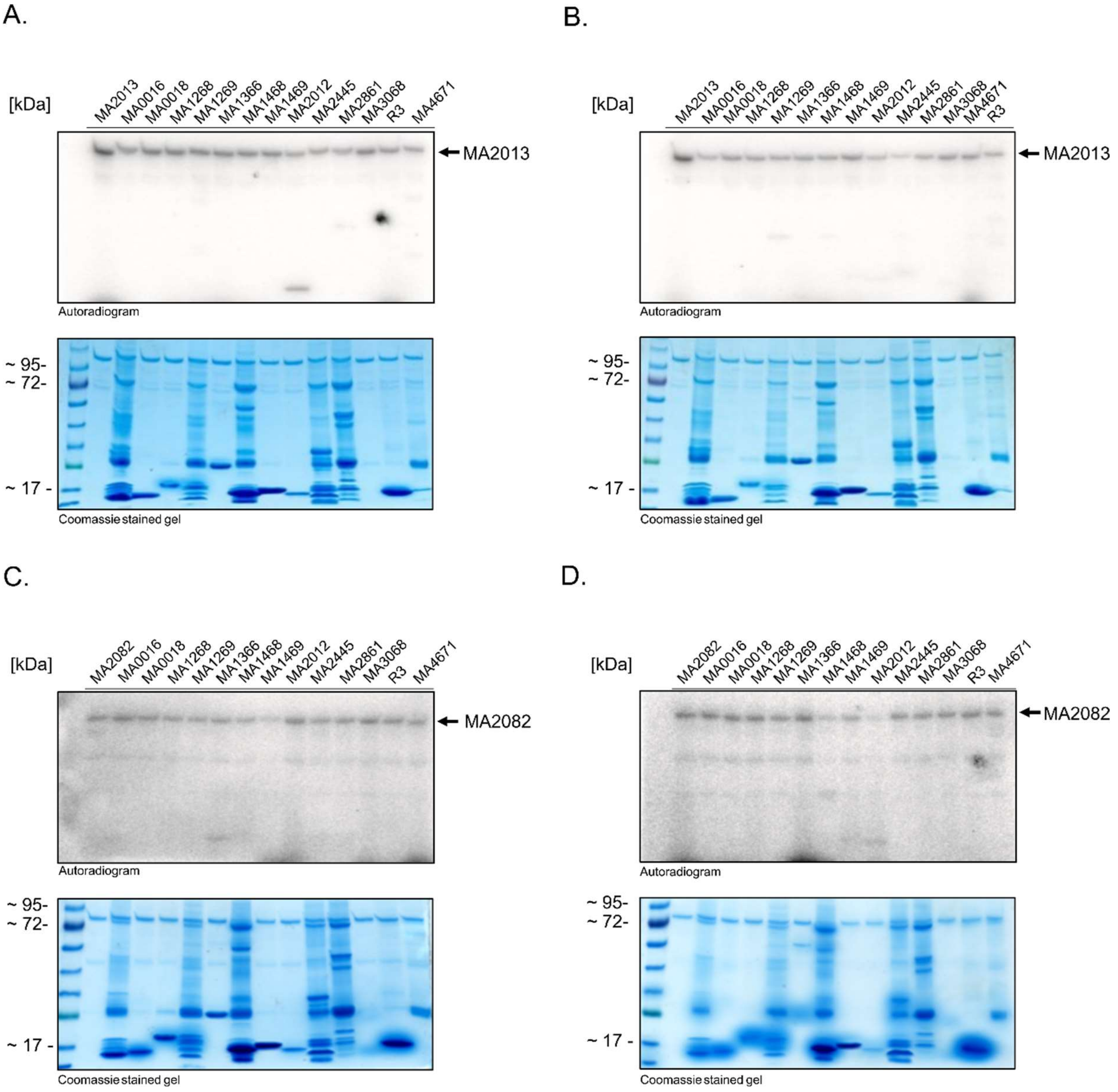


**Figure S8: Phosphotransfer profiling of MA2013 and MA2082.** The purified HKs MA2013 and MA2082 are incubated with radioactive labelled ATP and then incubated with each of the purified RRs. The reactions were analyzed by SDS-PAGE and phosphorimaging. (**A.**) Phosphoprofil of MA2013 after 30 seconds. Transfer is visible for the REC-only RR MA2012. (**B.**) Phosphoprofil after 60 minutes. Transfer of phosphate was not clearly identified, possible transfer to MA2445 and MA1268. A signal around 80 kDa corresponds to the autophosphorylated kinase, a second signal corresponds to the transphosphorylated RR. (**C.**) Phosphoprofil of MA2082 after 30 seconds. (**D.**) Phosphoprofil after 60 minutes. A signal around 80 kDa corresponds to the autophosphorylated kinase, a second signal corresponds to the transphosphorylated RR. Transfer is not clearly identified, MA1468, MA1469, MA2012 and MA1366 might be involved in the phosphorelay.

**Amino acid sequences of HK domains of putative HK and amino acid sequences of REC domains of RR of *M. acetivorans*.**

>MA3405
NMSHELKTPLNSIIGFSDLLKEEIAGPLNEKQSRYVQFISSSGKNLLEIINDILDLSKAESGEEDLNVEKFSVDESINKVISV
VLPQAQEKNIILNYQSENRTLWITADEGKFRQIMENLLSNAIKFTPAGGSIDVTLKQEGLLVTIEVKDTGIGIPEDSFEKIFK
PFIQIDSSLSRNFEGTGLGLTLVKKYVEMHGGNIYVESKIGEGSSFRFELPVTR
>MA2294
SMSHELRTPLNSIIGFSDMLLTQNFGPLNKKQLRYVNNISVSGNHLLKLINGILDLSKVEAGKMELKVEEFSVLDSISEVKVL
LTPLASKKDIQILSVVDKELTTIRADRTKFKQILYNLVDNAIKFTHEDGFVIVDARVEGDQAKIMVKDTGIGISKAGVKKIFQ
PFTQLENSEYGKQKGTGLGLSLVKKFVEMHTGKIWVESEFGEGSKFIFTIPLSL
>MA3962
NMSHELRTPLNSVIGFSDLLLEGAFGPLNTKQSKYVNNILISGKNLLEIINNLLDISRLEAGEKTLKYENVDIASLIGDVRMS
LLSPASVKRITVELKIDPSVGNVRADITKLRQILYNLVSNAIKFTPAKGKVVISACKKEGVLEVKVSDTGIGLSKDSHEKIFM
PFTQADSSAARGYGGAGLGLYIVRNFVDLHGGKIWVDSEVGKGSIFTFTLPAE
>MA1149
NMSHELRTPLNAVIGFSDLLLSETAGPLNEKQKRYTENISKSGSHLLDVINDVLDISRLELGNIELYYETVDIPGVIEEVRRV
LSSLSAEKNIRIEYKVEQGLKTIDVDRVKFKQILYNLLNNAIKFSSEDGKVNIKARSEGDMVEISVKDEGIGINEADYARVFL
PFVQIDESISRKHGGVGLGLALVKRFVELHGGQVWVEASPGKGSTFTFRIPKRP
>MA2555
NMSHELRTPLNSIIGFSDLLYEKIYGDLNEKQLKAVGNISRSGKHLLNLINDILDLSKVEAGKLELEYKEFELSSKLNSIKNL
LSPIADRKMIEVQIQVDESLNTIRADEARFAQIMYNLLDNAIKFSKENGLVEIDAKRKGDTVEITVKDYGIGIKVEDQSKLFK
PFSQIASFSSKKVQGTGLGLALVKQIVNLHGGYIWFNSRIGEGSTFAFTIPING
>MA1739
NISHELRTPLNSIIGFSDLLCEQIFGELNEKQLRYAGNISKSGKHLLSLINDILDLSKVEAGKMELDYTEFELAGKLNTIKNL
LAPIADRKSIQIEIEVDSRLTNLYADEAKFAQIMYNLVDNAIKFASENSPVNVGARMKGDKVEITVTDIGAGIKPEDQHKLFK
PFSQVDYFASKQHQGTGLGLYLVKQIVQMHRGYVWFRSVPGEGSTFAFAIPING
>MA3368
NMSHELRTPLNSIIGFSDMLYEQAYGELNRKQLRAIGNISSSGKHLLNLINGILDLSKIEANKMELNYREFDLATKLELIRNV
LYPVADKKNIDIEIDMDTELTKICADEEKFTRIMYNLVDNAIKFSYENSLVKIGARKNGNLVEITVTDAGIGIKAEDQHKLFK
PFSQVNSFPSKKFQGTGLGLSLVKQIVNLHGGYVWFRSEQGEGSTFAFAIPIAS
>MA2553
NMSHELRTPLNSIIGFSDLLYEKVYGELNLKQTKAVGNISNSGKHLLNLINEILDLSKVEAGSFELHYSTFWLAEVFAEVRDM
IFPFATSKGLKIELEIDSNLSRVYADKERILQVLSNLVTNAVKFSNENGCVKVKAVQMDGFLKITVADDGIGIAAADHEKLFK
PFSQIDSSFSKRYQGTGLGLALVKEIVQLHGGTVWFESEVGKGSVFGFSIPLPG
>MA4377
NMSHELRTPLNSIIGFSDILIERVFGELNEKQLKYVNNISGSGKHLLGLINDILDLSKVEAGKMDLHYSEFTVDSVFEEVKST
LFPLAQAKSLEINFVVGPDFGDIQADRSRLIQILYNLVSNAIKFTPEGGRVSVYCKKSGSRALFSVTDTGIGISSEDQKKLFQ
PFTQIDSSSARQYCGTGLGLALVKKIVNLHKGDIWVESELEKGSTFMFTIPLTK
>MA2348
NMSHELRTPLNAVIGFADILNEEGFGPLNKKQKRFVGNISTSGKHLLKLINEILDLSKIEAGKMEFKCTEFSVKEKFDEIKDI
LFPVFSKKKIRIEFEIGDGITTIYSDEGKFVQVLYNLVSNAIKFTEEGGFVKVSARKNGDMLQLSVKDTGIGITEEDLMKIFH
PFVQVDSFSTRQYEGTGLGLALVDQMVQLMGGEICVFSQPGIGSEFICMIPLKN
>MA0777
LKKAFYNLFDNARAHGDHVSEIDVSSHIVGESVVIEVKDNGIGVSPVMKELIFEKSVGRNTGLGLFLVRGILSITGMEITETG
IEGEGARFEIKVPPGN
>MA1957
TVSHELKTPLNSIIGFSDLLLEDSSGKLSERQARYINNISISGKHLLQFIDDILDLSKIEAGKTFLEPENFEFTKIFKDIEKV
FRPRVSGKKLSLNFEVDSGIKSFYADKMMFKQILYNLISNAVKFTPEEGSITVSAAKIGNMVRICVKDTGIGISREDMDSFFQ
PFKQPDSFFKRRYERTGLGLFLVKRFVEMHGGNIQAESVPGEGSSFIIELPLKT
>MA2256
TMSHELRTPLTAIIGFSELMLGEATGEFDELNRKFLGHISNSGKHLLSLINSVIDLSRIEAGKMDLEPDFFSLYDIFADTKSI
SSPLALKKNISMDFNVESDFLIYADRTRFKQIMYNLVSNAIKFTPAGGSVEVVGRRSENRIRVTVSDTGIGISQDEIKHLFKP
FKQINFALSREYESTGLGLVLSKNFVEMHGGRIWVESEPGKGSTFTFEVPVEI
>MA2013
NMSHEIRTPMNAVLGMLEMLLETSLTDEQREYLQLAHASAESLLSIIDDVLDFSKIEQNKLELEQISFELESLISHIINLLSG
KAGSKGLKLAFHIEKDLPSTFIGDPVRLKQVLFNILGNAIKFTKKGEIALSVEVYMPSERNSGLSNSDFPEEVALLFKVKDTG
IGIPPEKLSQIFDPFIQADASVTREYGGTGLGLAISSQLVELMEGRIWVESEVGKGCTFYFTVRVKR
>MA1270
VSSHDLQEPLRMIASYLQLLQRRYQGELDERADKYIYFAVDGASRMQSLINDLLEFSRVTTKAREFEPTDCESILNYVLSDLE
VSIKENEATVSYDSLPEIMADGTQLTQVFQNLISNAIKFRSKEAPKIHVSAEKEDDKWRFSVQDNGIGINPKYSEKIFEVFKR
LHKREEYPGTGIGLSICKKIIERHGGDIWVKSEPGRGSTFYFTLPASF
>RdmS
EDQQQKAIDTVINSSERLKHMVDSLLYLSLEQAGKIEYSFGEVEIKKILSDVYLNLVLLIDEKELKVEKELPASLPPIRGDKQ
KLTDLFTTLMGNSIKFTPHGGTLEVKAEEEEETIHITLKDSGTGIQKRLIPHLFHRIYQVDDSLTRRYQGLESGFYICKNIVN
AHEGEIWVESEEGSGTTMHVRLPKKK
>MsmS

EFVEEMMFPEKAEYGEIMDYETLYAIDSQQQKAVNTFIHYSEKLRRLVDSLLYQSLEKAGKIDYSFEETQLKDVLSDAFLNNV
FLIGEKALEVKKEVSASLSEIKGDREKLTALFTALIDHAIKFTPQGGKLALEVKEEAGNVHIVIADSGKGISKELIPYLFDRL
YQVNDSITRRYQGLESGLYICKNIVDAHKGEIWFESEEGLGNLMHVKLPK
>MA0203
EIHHRIKNNLQVISSLLDLQAEKFNSREDIKDSEVLEAFRESQDRVISMALIHEELYKGGEFDTLDFSSYIEELTENLFLTYR
LGNTDISLNMDLEENIFFDMDTAVPLGMIINELVSNSFKYAFQGRNRGEIRIKFRREENGKYINSGVKSKNGGCKSNNFTLTV
SDNGVGIPENFDIEDLDSLGFQLITSLVDQLDGKLELKRNNGTEFTVRFTVRD
>MA0490
EIHHRIKNNLQVISSLLSLEAEKFSDEKMLESFRESQNRVASMALIHEELYKGNELDTLDFAAYLQKLTADLFDSYNLGDSGI
SLKLDLEKIHLDMDIAIPLGIIVNELVSNSLKHAFSAGKAGEIHISFCKKESFAANDDIPGPCPFCTGKNNLHYILTVADNGK
GIPEEIKFPNTDSLGLQLVNLLVEQIDGYIELKSDSGTKFTIWF
>MA0551
EIHHRIKNNLQVISSLLDLQAEKFQNKEVLEAFRESQNRVTSMSLIHEELYKGGENNTLNFSTYLQKLAENLFQTYSLKSKKV
LLCMDLEENTLFDMDIAVPLGIIVNELVSNSLKHAFTEEEEGEIRIKLCREEKGNEMHRSLFSLSISDNGKGIPESTKLESIE
SLGLQLVSILVDQLDGKIELKRTHGTEFRITFNVTE
>MA0552
EIHHRIKNNLQVISSLLDLQAEKFNNREDIKDSEILEAFRESQDRVISMALIHEELYKGGGFDTLNFSSYIEELAENLFQTYS
LGKADISLKMDLEERIFFDMDIAVPLGIIVNELVSNSLKHAFTEGEEGEIRIRLCREKKNNELDKSLFSLTISDNGNGIPENV
ELKSFESLGLQLVSILVDQLDGKIELKQEQGTKFRITFKVAE
>MA0619
EIHHRIKNNLQVISSLLSLEAEKFSDERTLEAFRESQNRVVSMALIHEELYEGKGMDTIDFAVYLRKLTLDLFNSYTVETRKV
RLNLDLEQVYLGMDTAIPLGIIVNELVSNSLKHAFPSGGEDEIRINLRRMEDSTSKPERVPGCQNGKVFHYMLTVTDNGRGFP
EEINFQNSESLGLQLVNILVEQIDGCIELKRDRGSEFAIFFDNPG
>MA0620
EIHHRIKNNLQVISSLLSLQAEKFEDREVIEAFRESQNRVASIAMIHEELHGGENLDSLDFADYLQKLTADLFDSYRVGKEGV
SLKLDLENVYLGMDTAIPLGIIVNELVSNSLKHAFSDKNEGEISINLHSNDASHSLVELSCSEPEFLENDDFHYTLTVADNGR
GIPEKMDFRNTDSLGLQLITILVEQIGGCIELNRDYGTEFVINFGNSE
>MA0758
EIHHRIKNNLQVISSLLDLQAEKFRDKDVLEAFRESQSRVLSMSLIHEELYKGGETDTLDFSTYLEKLAENLFRTYSFRSKNI
CLSMNLEENAFFNMDIAVPLGIIVNELVSNSLKHAFTEKEGDIRIRLCREESDCEMSKSLFSLTISDNGKGIPENVELESVES
LGLQLVNILVDQLDGNIKLKHDQGTECRIEFKVME
>MA0759
EIHHRIKNNLQVISSLLDLQAEKFRDKEVLEAFRESQSRVLSMSLIHEELYKGEGTDTLDFPTYLQKLAENLFQTYSFRSKNI
RLYMDLEENAFFNMDIAVPLGIIVNELVSNSLKHAFTENKEGEIRIKLCREEKNNEMQESIFSLTISDDGKGIPENIELENIE
SLGLQLVNILVDQLEGNIELKRDQGMESRIEFKVMER
>MA0970
EIHHRIKNNLQVISSLLDLQAEKFRGKKNIEDSKILEAFKESQDRVISMALIHEELHKSGEIDTLNFSAYIHELSGNLFLSYR
LGNDGISLDMDIEEDIFFDMDTSVPLGMIVNELVSNSLKHAFSDRDKGEIRIKLHRKKNRESDIEDCCIAFILSVSDNGIGIP
KDLEIEDIESLGLQLVTTLIDQLDGELELKRDDGTEFIVRFSITE
>MA1274
EIHHRIKNNLQVISSLLDLQAEKFRSREHVEDSEVLNAFKESQERVISIALIHEELHEGKGTDTLNFSPYLQRLVKNLFQIYN
LGNVDISLSMDIEENVFFDMDTAVPLGLIVNELVSNSLKYAFKGRDKGVIRIKLSREGNGEIMSNREESKKEGHEDTNFVLTV
SDNGVSIPEDFNLENSDTLGIQLVTTLVDQLDGRLEMKRGSGIEFIVRFPITE
>MA1322
EIHHRIKNNLQVISSLLDLQAEKFGNKKYIMNSEVMDAFRESQDRVISMALIHEELHKSEGLDTLNFSPYIEELAENLFQTYR
LGNSNICLNMDLEENIFLNMDTAVPLGIIINELVSNSFKHAFTEDEEGEIRIKLHREENGEHKKERNKSTDFVLTVSDNGAGI
PENLDIEDLGSLGLQLVTSLVDQLDGELELKRENGIEFTIRFTVTE
>MA1470
EIHHRIKNNLQVISSLLELQAEKFDNLEVLEAFRESQNRVATMAIIHEELYRSRNNETLDFSAYLQKLTADLFHSYLVRKGDV
GMQLNIEEIFLGMDTAVPLGIIINELVSNSLKHAFPSGRKGEIYISLCRTDENNENKSISNIINNMDAKSPVDKNIQYMLVIS
DNGLGFPENVDFTNTSSLGLQLVNILVEQLEGAIELENDSGTKFKIWFKEPC
>MA1627
EIHHRIKNNLQVISSLLDLQADKFDNPKVIEAFRESQNRVISMALIHEELYKGEGNDTLNFSTYIKELAGNLFQTYSLTSKNI
CLCMDMEKNVLLNMDTAVPLGIIVNELVSNSLKHAFSGKEGGEIRIKLRRKGNGSRKKEGDKATSFILIVSDNGIGIPENLNI
QDVDSLGMQLINTLVDQLDGKLELKRGNGTEFTIKFAVAE
>MA1628
EIHHRIKNNLQIISSLLDLQAEMFKGRSNIRDSEVLKAFAVSMDRVLSIALIHEELYKGKNIDVLNFSEYIKKLADNLLITYR
LETDVNLNLDLEENLFLNMDAAIPLGIIINELVSNSLKYAFPDRDKGEIRIKLRREEKGECKINGCKSADFVLTVSDDGIGIP
ENLDIKDLDSLGLQLVISLVDQLDGELKLKRGNGTEFTIKFAVTE
>MA1630
EIHHRIKNNLQIISSLLDLQAEKFEDKNVTEAFREGQNRVISMSLIHEELYKGEGTDTLDFSVYLKKLAENLFQTYNLSSKNI
NLSMDLEKDTFLDMDTAVPLGIIVNELVSNSLKHAFIEEGKVRINLCREERNYDTNESLFSLTISDNGKGMPEDLELESAESL
GLQLVNILVDQLDGELELKRAQGTEFTIRFKVVE
>MA1645
EIHHRIKNNLQVISSLLDLQAEQFKNRENIKDSEVLEAFRESQDRVISMALIHEELYKGGGFETLNFSPYIKELVENLFQTYR
LGDIDISLNIDLEENVFFDMDTAVPLGMIVNELVSNSFKHAFIGRDKGEIRIELYREESAEFESENRKSTNFILTVSDNGVGI
PDNLDIEDLGSLGMQLVVSLIDQMNGELELKRNNGTEFTMKFTVTE
>MA1646

```
ELHHRIKNNLQVISSLLDLQADLFKGKKTITDSEVLKAFNESIDRVLSIALVHEELYKGKNIDLLNFSQYIKELANNLLLTYS
LKTDVSLNFDLEENFFLDMDTAIPLGMIINELVSNSFKYAFPERDKGEIRIKLRREEKGKCKINGCKYADFVLTVSDDGTGIP
ENLNVKDLNSLGFQLVTSLVDQLGGEFELKRNNGTEFTMGFSVIE
>MA1704
EINHRIKNNLQVISSLLSFEAEKSTDPEILEAFRETQNRIASMSLIHQELCIGETYTIDLADYLRKLTAGIFSSYLVGNERIN
LRLDLEQVYMETDTAVPLGIIVNELVSNALKHAFLPGKEGEIRVNLSRMKNCEKEFKNSRSSGIVPGYSNGKNLQFILTIEDN
GRGIPELGDHQNKDSLGLQLVNIFVEQIGGSIERKKDKGTKFNIRFNKLE
>MA1844
EIHHRIKNNLQVISSLLELQADKFKDREVIEAFRESQNRVASMAIIHEELYRAGDIETLDFSAYLRKLTSDLLSSYTVRKEDV
KLKLEAEDTFLGMDTAIPLGIIINELVSNALKYAFPAGRRGEIRIKLCRKEINENKNIDEIISNNYRGSSIKNSYLYSLVVSD
NGLGFPENVDFKNTDSLGLQLVNILVEQLEGTIEMEKNGGTTFRISFTETE
>MA1878
EIHHRIKNNLQVISSLLDLQIDIFSNREICKTPEVIEAFRESQNRVVSVALIHEELYKSKGMDSLDFAAYLQKLTKNFLKSYN
IDADDINLKLDFEQVYLGMDTAVPLGIIVNELFSNSLKHAFPNKREREIKITLQKEENVYKKASFHYMLTVTDNGKGIPEEID
IQTADSLGLQLVNILVEQIDGCIELKRNKGTEFTIWFNNIE
>MA1991
EIHHRIKNNLQVISSLLSLQAEKFRDQEVLEAFRESQDRIISMTLIHEELYKGGGTDTLNFSKYIQKLTENLFRIYGLKSKNI
CLFMDLEENAFFNMDTAVPLGIIVNELVSNSLKYAFIDKQEGEIRICLCREEKNDGMNRSLFSLIISDNGTGIPENVELENVE
SLGLQLVNILIDQLDGEIELKRDHGTEFRINFEVAE
>MA2082
EIHHRIKNNLQVISSLLDLQAEKFRDKEVLEAFRESQNRVVSMSLIHEELYKGEGTDALDFSAYLRKLSEKLFQTYSLSSKNI
RMYMNLEENTSFNMDIAVPLGIIVNELVSNAFKHAFPEKVGEIRIQLQKEKMCNEVNRSLFSLIISDNGIGIPEGVELASFES
LGMKLVNTLVDQLGGKIEIIRAHGTEFRITFNVTE
>MA2266
EIHHRIKNNLQVISSLLSLQAEYFSDPKVKESFKDSQNRVISMSLIHEELYKTRETADIETFDFKVYIQKLATELFKSYLVGS
EDIRLKLDVESAFLGMDTGIPLGIIVNELVSNSLKHAFPEGRSGEIQIKLHRTGSSQKKGCNKHPGNCEISEFLLTVSDNGIG
FPEDLDLKHTSSLGLQLVNILVEQIEGSIELERGKGTEFRLKFREQE
>MA2732
EIHHRIKNNLQVISSLLDLECDSLLSGTPDHKKIAEAFRESHNRIISMSVIHEELYNSRDMETINFASYLKKLTDDLFKSYKV
GNSDIKLYLDVEDFFFEMDNAIPLGIIVNELVSNSLKYAFPDGRNGEIHIELQALDDKKSSNTTADFSMQNNVGPSSYFRLTV
GDNGIGFPGSFDFKNISSLGLQLVNTLVDQIGGSIEIGNGPGTKYNILFKDS
>MA2757
EIHHRIKNNLQVISSLLDLQAEQFKNRECIKNSEVLEAFRESQARVISMALIHEELYKGDGLEMLNFSPYIEELAKSLFHTYR
IGNSDIRLKLDLEQNISFDMDTAVPLGIIVNELVSNSLKHAFPDGGTGEITIKLHREENGEQINNPNKNCSVNFILSVSDNGV
GIAENLNIEDLDSLGLQLVTTLVEQLNAELELKRNNGTEFILKFIVTE
>MA2784
EIHHRIKNNLQVISSLLDLQAEKFKDREDIKDSEVLEAFRESQDRVISMALIHEELHRNEGLDKLNFSQYIKELADNLFLTYK
LGNDGTRFNKDIEENIFFDMDTSVPLGIIINELISNSLKYAFQGRNHGEIQVKLYREEDREQDIEDLNSTAYVLSVSDDGVGI
LKDLDIEELDSLGLQLVTSLVKQLNGELELERNNGTEFIIRFTVTE
>MA3346
EIHHRIKNNLQVISSLLDLQAENFSKHEVCKTPKVVEAFKESQDRVISIALIHEELHENGETDTLDFSPYLEKLVDALFQTYR
LGNARITLKKELEKNIFFDMDVAVPLGLIVNELVSNSLKHAFSDRDSGEIDIKLCREISPEQETDLCSENETKSRRETGFILT
VSDDGTGISDAVDLENSDSLGLQLVKILVDQLEGEMEVKREKGTEFTIRISVAE
>MA3370
EIHHRIKNNLQVISSLLDLQAGKFNNKEHIRDSEVLEAFKESQDRVTSIALIHEELHEEEGKTTDTLNFPIYLQRLVKNLFRT
YTLGNIDISLNLDLKENIFFDMDTAVPLGIIVNELVSNSLKHAFSGRNKGIIQIKLFSEEAGNEPNSKRKLSMEKLIKEKLAE
EELAEEELAKEELAKERIPGKSTGYTLIVSDDGIGIPEEIDIKNPETLGLQLVNILVDQLDGKIKLEREHGTEFIINFSVEE
>MA3481
EIHHRIKNNLQVISSMLSLQAEKFSDEETLEAFRESQNRVTSMALIHEELYEGKDMETLDFAVYLRRLIGDLLSSYTVGNREI
DLKLDLKQIYLGMDTAVPLGIVVNELVSNSLKHAFPAGKRGEIRISLSRSENYEKRHENGNFEVERYIETGSRSIEPDEMGIE
KSEEGIVRCSGSKRGIEKSEKDPVFMLIVADNGNGFPQNIDFRNTDSLGLQLVSILVEQINGSIELNRDEGTEFRILFGKVG
>MA3543
EIHHRIKNNLQVISSLLDLQAEKFNKREGIKDSEVMEAFRESQDRVISMALIHEELHKSGGLDKLDFSSYIKELADNLFLTYR
LGTIDVSLNMDLEKNIFFDMDTAVPLGMIVNELVSNSLKHAFLGRDRGEIRIKLRREGNRECRNSIGCVESNSKDCESPSFTM
IVSDNGVGIPKDLNIEELDSLGFQLVISLVEQLDGELELKRDKGTEFIMKFTVTE
>MA4026
EIHHRIKNNLQVISSLLDLQAEKFSHREAVPTLEILEAFKESQNRVISMSLIHEELYKGEGTDTLNFSVYLRKLAENLFQTYS
LCSKNIRLYMDLEENTFFNIDVAVPLGIIVNELVSNSLKHAFAENKEGEIQIKLCREESDCEMSKSLFSLIISDNGKGIPENV
ELGSVDSLGLQLVSILVNQVDGKIELKKAQGTEFRIIFSITE
>MA1267
EIHHRIKNNLQVISSLLDLQAEKFEDPTIRQAFRESQNRVISMALIHEELYESGEIGTLNFAAYMQKLVENIFECYNIGDHKA
HLHLEIEEKTFLDMDTAVPLGIIVNELFSNSLKHAFPERDEGIVKIRLCREENWECKSNSVENTGVENPGRDNTRGDNKSINL
ILTISDNGVGMPETVDMESPCTLGLQLVTILVDQLDGEIELKRDSGTEFIIKIAVKQ
>MA1463
EINHRIKNNLQIVSSLLDLQAEQFSDKKVIEAFRESENRIVSMSLIHEELYESGNLDILDFSSYIRKLIDDLCRSYSTESSHI
RIKLDVETVFLGVNLAVSLGIIINELFTNSLKYAFSPGKGEEISISLLREDTRGQDQRPEAISEKVFLTASEDSENFTLVFAD
NGKGFPEESQFQKHGIPWAAACKCSCEPDRGEHRP
>MA2890
```

EIHHRIKNNLQIVSSLLSLQADKFKDKDVIEAFRESENRVISMSIIHEELHKSEDTTNIDFAAYLRKLTSELLYSYKVGNEKV
RFFLDVDNVFLGIDTAIPLGIIINELFSNSLKYAFLKSAEGEIRISLYRQPEVLELCSTSGRGQNTGSVVHSSFTLVYSDNGG
RFPENIDFKNPETLGLQLVNALVEQLDGTIELEKGGETKFIIRFDDKGLPGKS
>MA0014
RISTEQLDKLMNLVGELVINRSRVKELTGESKSKDLELALSEFQKLTRELQEEVLEIRMVPLDHITNIFPRMIRNLAREQNKK
INLVIRGKEIKLDRAIMEEIGDPLVHLLRNAVDHGIELPEQRVELGKEETGTIMITASKQQNYVLVKIEDDGRGINAKEILQA
ALEKGFISRDEAEQLSEREAIQLIFAPGFTTASTVTDLSGRGVGMDVVKNRIEHLGGSVKVESKLGFCSRFELRLPITI
>MA3066
GSSHHFSESAASAKTPLETQRQETIRVRTSNLDNIMNLVGELVINKGRLLQISQEYNLPELEEATGALDKSISSLQDEVMLIR
MVKIERVFSKFPRMVRDLSRKFDKNIEFEIEGQDTELDRTILDEISDPLVHLVRNAVDHGIEYPEAREKAGKNEVGNIKLSAR
REKNNVIIEIEDDGKGIDVEVLKKKAVEKGIFSAFDVENLSEEEIRMIIFSLGFSTKESPTEISGRGVGMDAVKTAVEKLGGK
VRVYSKKGEFTRIRIDLPPTV

>MA0016
MARVMIVDDAEFMRMVIRDILLKHGHEVVAEVGDGEEAIQTYLEVKPDLVLMDIIMPDMDGKEALQKLLLIDPDAKVVMCSSL
GQQALITESMKIGAMGFIIKPFEPDGMLDVIKKIAEPN
>MA0018
MPEILIVEDNLLNLVIEADLLKSCGYDPKKAKNGFEALEVLSKVKVDLVLMDMELPKMHGLELLQRIKCNPETQGIRVVAVTG
HCDPESEQEFLKAGCHAVLSKPINFDLFGAQVKEFLTATNSPG
>MA1268
MKTKVAAKPIEILLVEDSEGDVGLIEEVFEEAKIRNNLHIVEDGEEAILFLRGEKQFSGISRPDIILLDLNLPKKDGREVLEE
IKEDDDLKNIPVVVLTTSKAEEDVLKSYNLHANAYVTKPVDFDQFIRVIKSIEDFWLEVVKLPSK
>MA1269
METWTAFKPVDILLVEDNKGDVGLIEEVFESSKVRNKLYVVEDGEEAVHFLLREGKFSDVPRPGIILLDLNLPKKDGREVLEE
IKEDDDLKNIPVVVLTTSRAEEDILESYKLHANAYVTKPVDFDQFIKVIKSIENFWLEVVNLT
>MA1366
KILIMGNGNNVHNNLQKVLEAENYNVVSASDNFSAIETVNEEKPDLVLLDTVYLETDGFEICRQLKDSPRYWWIPIMMLSERN
KTEDGIKAFDSGADDYITMPFNPLELKARVGMIL
>MA1468
MDKAKILVVEDQNIVALNLRNRLKNMGYIVPTIAISGEEAIRKTELTTPDLVLMDIMLKGDMDGIEAARIIKSRFSAPVIYLT
ACTDIGILERAKLTEPAGYISKPFKEKDLVSNIEVALQKNKLGKVTEE
>MA1469
MKKAKILVVEDQNIVALNIRNKLKNLGYTVPGTASTGEEAIRKAELTNADLVLMDIMLKGDMDGIEAAREIKARLKIPVLYLT
AYTDDETLERAKMTEPAGYISKPFKEEDLHSNIEMALHKHRTEKKEIENSDSDE
>MA2012
KVLIAEDEPISNLWLKNTLTRWGYEAISTRDGYEAWEVLNESDFPRVVILDWEMPKMKGIEVCEKIKKDPRLSSIYIILITGR
DLTEDMEAGFKAGADDYLKKPFDNRKLKTKLDTAR
>MA2013-R1
NVLFAEDHPINQKLILGLLEKKGHKLTLVTSGKDALDALSRRDFDAVLMDIQMPGMDGLEATRRIRDPSSGVRRHNIPIIAFT
ARALKEDREKCFEAGMNYYISKPLKKEKLLNILEDIR
>MA2445
KVLIVDDKKENVELMEAYLAVEPYDVITAYGGKEAFQKVKEEKPDIILLDVMMPEVNGYEVCKILKGNPETQFIPVLMLTALS
ELEDRIRGIEVGADDFLTKPINRLELKTRVKSLL
>MA2861
MKVLLVDDDPVFLELSKTFLEVFHDINSDTVESARQALEKLDELSYDVVVSDYDMPYMDGISFLKTIRDKRINIPFILFTGVG
KEEIKSQAIENGVDSLIQKRGDPKAQYSELSKRIWQIVKNGSG
>MA3068
MAKVLIVDDTAFMRKLLKNILFGAGFDIAGEAENGKQAVEMYKGLKPDVVTMDVVMPEMTGIDALKQIKALDKDAKIVMCTAI
GQENIVKTAIKLGARGYIIKPFQAPKVIEEIKKVIGA
>MA4376
MKEILIVEDNPMNMELILDLLEFYGHRVTEAEDGIKALERLAEKKFDIILLDMQLPKMDGLEVLDRIKKNPATADIPVIAVTA
HAMKGSEEHFIEMGCVDYISKPIDIHRFRSLIDKYLG
E
>MA4377-R1
LVLVVDDDINSNELISVVLREAGYSTASLHNGKDVLEVAKKLKPDVITLDVLLPDTSGWNVLKQLKSDLDTTSIPVLIISVTD
NNELGVALGATYSFTKPVRRVELLDSLREIT
>MA4377-R2
KVLIIDDDENAVELLSSMIESEGFEIVKAYSGQAGLDKLFSEQQPDILILDLLMPEISGFEIISRLRDGEQTKDIPLIVCTAG
EFTEKNIEKLNGELKGHLISIMKKGTFGRKELINRIKQLA
>MA4671
MEMKSILVVEDSPVILELISFFLTSSGYESRETGDGFDALKIAEENRFDLILLDKQLPGFDGLEVLKKIKKIFEIRKTSVIAL
MAHAMQGDEDRFLKAGCNGYISKPIDIDRFKLILDTCTGGYQVL
